# A *Mycobacterium tuberculosis* rare variable antigen vaccine reduces lung pathology without affecting bacterial burdens

**DOI:** 10.1101/2025.08.01.668220

**Authors:** Zachary P. Howard, Alexander Mohapatra, Weihao Zheng, Joel D. Ernst

## Abstract

*Mycobacterium tuberculosis* is the leading cause of death globally due to a single infectious agent. Currently, no reliable vaccine against pulmonary tuberculosis, the primary adult disease caused by Mtb infection, exists. CD4 T cells are essential in protection against Mtb infection and inducing a protective CD4 T cell response remains the goal of most Mtb vaccines currently in testing. However, most Mtb T cell antigens do not exhibit antigenic variation, suggesting that T cell recognition does not drive selection of escape mutants. We utilized a set of antigens that do exhibit sequence diversity within human T cell epitopes and tested the impact of vaccination with these rare variable antigens (RVMA) using a DNA vaccine platform. We found vaccination with RVMA significantly alters the immune response to Mtb infection in both C57BL/6 and hypersusceptible SP140^-/-^ mice without reducing bacterial burdens. RVMA vaccination of hypersusceptible SP140^-/-^ animals prevented necrosis and altered the lesion composition reducing tissue damage and increasing CD4 T cell distribution. Reductions in pathology were associated with increases in RORγt-expressing CD4 T cells and decreases in monocyte-derived cells in the lungs prior to the development of necrotic lesions. These results suggest T cell responses to certain antigens may be involved in preventing pathology without significantly changing bacterial burdens.

**Significance:** Mtb is the leading cause of death worldwide due to an infectious agent, a title it has held for most of human history. Despite significant efforts, an effective vaccine that prevents the morbidity associated with pulmonary tuberculosis, the most prevalent form of adult TB disease, has yet to be developed. A lack of correlates of protection and ideal antigen targets have hampered the development of a subunit vaccine. Mtb is unique in that the CD4 T cell epitope sequences of Mtb are predominantly under purifying selection. Targeting antigens that are an exception to this T cell epitope sequence conservation can inform our understanding of what the most optimal antigens are for inclusion in an effective vaccine against Mtb. Our study describes the effects of a vaccine consisting solely of antigens that exhibit epitope sequence diversity on the outcome of infection and identifies possible benefits to lung pathology. Given the ongoing interest in prevention of disease (POD) vaccines for TB, these results can inform future vaccine design and the search for correlates of protection.

## Introduction

*Mycobacterium tuberculosis* (Mtb) has returned as the greatest cause of human mortality due to a single infectious agent [^1^]. Despite discovering Mtb as the causative agent of tuberculosis (TB) over 140 years ago, we still lack an effective vaccine to prevent pulmonary TB in adults [^2–5^]. While BCG remains the only approved vaccine against TB, it has variable efficacy in preventing TB disease [^6–8^]. Recent attempts to develop an effective subunit vaccine have yielded mixed results, with only one candidate, M72A/AS01E, providing approximately 50% efficacy in preventing TB disease [^9–11^].

Most subunit vaccine candidates aim to generate a protective Th1 CD4 T cell response, and the evidence supports the notion that CD4 T cells provide protection against TB [^12–15^]. However, the optimal antigens to target with a subunit vaccine remain unknown. Indeed, antigen selection was a primary difference between an unsuccessful vaccine and an efficacious one [^9,10^]. Interestingly, Mtb CD4 T cell epitope sequences are uniquely conserved [^16,17^]. While other bacterial pathogens utilize antigenic variation as a mechanism of immune evasion, Mtb is an exception [^18^]. In fact, hyperconservation suggests Mtb CD4 T cell antigens may be under purifying selection due to a positive effect of T cell recognition on the bacteria, such as by promoting transmission. Therefore, we hypothesized that conserved Mtb T cell antigens may be suboptimal and that the antigens that exhibit sequence diversity within epitope sequences are alternative targets for vaccination. We previously identified a small group of antigens that exhibited sequence diversity within CD4 T cell epitopes, across strains and lineages within the *Mycobacterium tuberculosis* complex (MTBC) that we hypothesize are under diversifying selection due to CD4 T cell responses that are more detrimental to the bacteria than are T cell responses to conserved antigens [^17^].

Lung pathology is an important aspect of TB [^12^]. As a human-specific pathogen, Mtb relies on human-to-human transmission via the aerosol route. Humans with more extensive lung pathology shed a higher number of bacteria and are more infectious [^19–22^]. One form of TB lung pathology termed cavitary TB is associated with enhanced transmission and is the development of cavitary TB is facilitated by CD4 T cells, since individuals whose CD4 T cells are depleted by HIV infection rarely exhibit cavitary TB [^23^]. Moreover, in people infected with HIV that have active TB, the frequency of cavitary TB is inversely proportional to the peripheral CD4 T cell count [^21^]. Together, these results indicate that some CD4 T cell responses can be detrimental in humans with TB.

Given the significance of lung pathology to TB morbidity and transmission, recent vaccine trials have focused on prevention of disease [^10,12^]. While many studies have demonstrated the protection afforded by CD4 T cells that make IFNγ, the fact remains that those same CD4 T cell responses can be associated with more cavitary TB and higher bacterial shedding [^19,21^]. In mice and humans, the CD4 T cell response can also be more pathogenic if immunoregulation is lost, such as in the case of anti-PD-1 therapy or genetic deficiency of PD-1 and other immunoregulatory factors [^24–27^]. Most mouse studies of Mtb infection use C57BL/6 mice, a resistant strain that exhibits limited lung pathology. More highly susceptible strains of mice, such as C3HeB/FeJ mice develop more lung pathology, but genetic differences from the C57BL/6 mice make it less tractable for in-depth immunology studies [^28^]. Recently, an isogenic C57BL/6 mouse strain was developed by disrupting the SP140 gene to generate a susceptible mouse model that exhibits severe lung pathology similar to that in C3HeB/FeJ mice [^29^]. Using this mouse strain, we evaluated the ability of a vaccine targeting Mtb rare variable antigens (RVMA) in preventing severe lung pathology while retaining the ability to evaluate T cell responses in a C57BL/6 genetic background.

In this study, we report on a vaccine containing antigens selected based on increased sequence variation and found that this vaccine reduced necrosis in the hypersusceptible SP140^-/-^ mouse strain without significantly reducing bacterial burdens in the lungs or spleen. We identified differences in the cellular response to infection in vaccinated animals that is associated with reduced tissue necrosis. We also characterize the lesion structure and composition in vaccinated animals, demonstrating a shift from necrotic to non-necrotic fibrotic lesions.

## Results

### Variable antigens are immunogenic and antigenic in C57BL/6 mice

A DNA cassette consisting of a fusion of 4 RVMA [^17^] was synthesized and cloned into the commercially available pVAX1 DNA vaccine vector backbone to create pCas4 (Fig. 1A). To assess immunogenicity of this vaccine plasmid, C57BL/6 mice were vaccinated 3 times, 3 weeks apart and IFNγ responses to the pCas4 antigens were assessed 3 weeks after the final vaccination. Spleen cells from vaccinated animals were stimulated with pooled overlapping peptides spanning the 4 RVMA and IFNγ was measured after 72 hours of stimulation. IFNγ was detected in response to 3 of the 4 RVMA, with RimJ inducing the highest magnitude responses (Fig. 1B). Given that these candidate antigens were identified in clinical isolates from patients and epitopes were confirmed to produce T cell responses in Mtb-exposed patient samples [^17,30^], we next sought to determine the antigenicity of the 4 RVMA by determining the magnitude of CD4 T cell responses during infection in mice. We performed ICS on lung cells from Mtb H37Rv-infected C57BL/6 mice that were stimulated in vitro with pooled overlapping peptides spanning the pCas4 antigens as well as an Mtb whole cell lysate or the immunodominant antigens Ag85B and ESAT-6. IFNγ-producing CD4 T cell responses were observed for the same 3 antigens that induced a response in vaccinated animals Rv0010c, Rv2719c, and RimJ (Fig. 1C). However, the magnitude of the IFNγ response was lower for all pCas4 antigens as compared to the response to the conserved immunodominant antigens Ag85B and ESAT-6 (Fig. 1C). Subdominant antigens have been shown to provide protection when incorporated into vaccines [^31^], which indicated that responses to these antigens could still prove effective as vaccine antigens. We therefore determined the impact of vaccination on the magnitude of CD4 T cell responses during infection by challenging vaccinated animals with the matched Mtb Erdman strain at 28 days post-infection. Vaccinated animals had a significant increase in the IFNγ^+^TNF^+^ CD4 T cell responses to Rv0010c and RimJ, with a trend towards increased responsiveness to Rv2719c, as compared to mock-vaccinated animals (Fig. 1D). Vaccination with pCas4 also enhanced the CD4 T cell response to the WCL, which indicates the enhanced T cell response may not be limited to pCas4-specific T cells (Fig. 1D). This IFNγ^+^TNF^+^ CD4 T cell response has been associated with significant reduction in bacterial burden in other Mtb vaccine studies, which suggested pCas4 could be efficacious as well [^32^].

**Figure 1.**
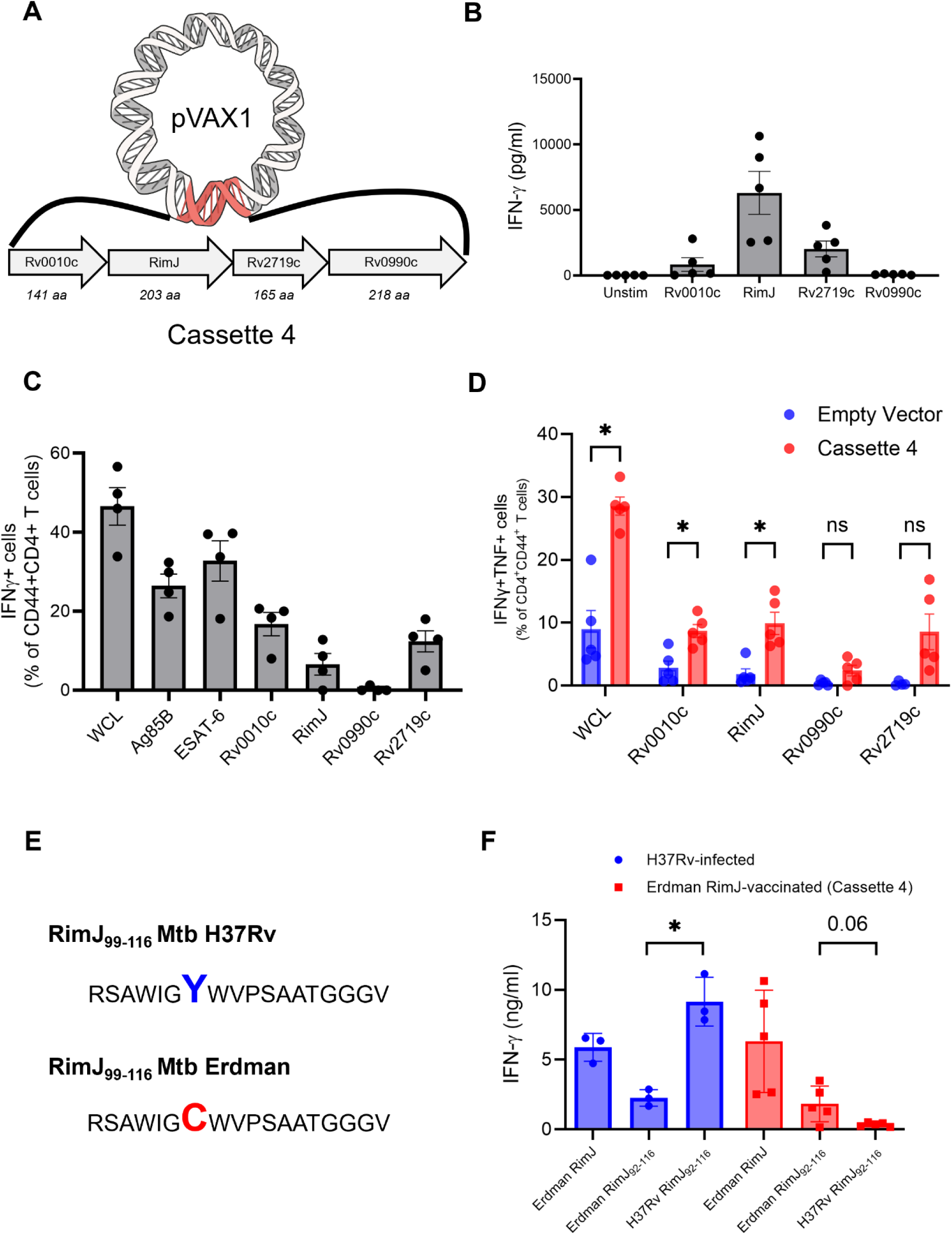
Immunogenicity of a DNA vaccine encoding 4 variable antigens (Cassette 4). Rare variable antigens identified previously, were selected and the Mtb Erdman sequence was cloned into the commercially available DNA vaccine vector, pVAX1. **A)** Depiction of the pVAX1 vector containing the 4 chosen antigens used in Cassette 4. **B)** IFNγ concentrations in culture supernatants from Cassette 4-vaccianted splenocytes (n = 5 mice) stimulated with the vaccine antigens or unstimulated (unstim) for 72 hours. **C)** IFNγ^+^ CD4 T cell frequencies from lung leukocyte cultures from Mtb Erdman-infected mice (28 days post-infection). Cells were stimulated with pooled peptides spanning the indicated antigens overnight or an Mtb H37Rv whole cell lysate (WCL) in the presence of GolgiStop and GolgiPlug. All frequencies have the unstimulated cytokine positive frequency subtracted (background subtracted) (n = 4 mice). **D)** IFNγ^+^TNF^+^ CD4 T cell frequencies from lung leukocyte cultures from Mtb Erdman challenged Cassette 4-vaccinated mice at 28 days post-infection. **E)** Depiction of an amino acid substitution in the Mtb Erdman strain within the primary predicted epitope of the RimJ antigen. **F)** IFNγ concentrations in culture supernatants from Mtb H37Rv-infected mouse splenocytes (28 days post-infection) and Cassette 4-vaccinated mouse splenocytes stimulated with a peptide pool spanning the Erdman RimJ protein or single peptides corresponding to the H37Rv or Erdman epitope sequence. At least 3 mice were used for each experiment. Welch’s t tests were used to compare groups, and significant differences are indicated by a * (p-value < 0.05).

As the antigens in pCas4 were identified by their sequence diversity across strains of Mtb, we assessed protein sequence differences between the two strains of Mtb used in this study, H37Rv and Erdman. Despite both being related lineage 4 strains, we identified multiple amino acid substitutions in the pCas4 RVMA, RimJ. One of these substitutions was located within the predicted core epitope for the mouse H2-IA^b^ allele, RimJ_99-116_ (Fig. 1E). The substitution, Y105C, in the Erdman sequence induced a lower response in spleen cells from H37Rv-infected mice as compared to the H37Rv-matched RimJ_99-116_ sequence (Fig. 1F). In mice that were vaccinated with pCas4, which was designed using the Erdman sequences, the Erdman RimJ_99-116_ peptide induced greater responses than the H37Rv-matched peptide but did not match the total response induced by the pooled overlapping peptides (Fig. 1F). We further determined a secondary epitope in Erdman RimJ, RimJ_182-197_, that likely accounts for this difference (Fig. S2). While we were excited to discover a naturally occurring substitution in one of the pCas4 antigens that altered the CD4 T cell response, we also recognized the possible impact on vaccine efficacy if the vaccine antigens and Mtb antigens were mismatched. Due to the nature of the sequence variability of the pCas4 antigens, we used the matched Erdman strain for all further challenge experiments.

### Variable antigen vaccination reduces lymphoid aggregates during chronic Mtb infection

Recent vaccine efforts have focused on prevention of disease rather than prevention of infection [^10^]. Therefore, we determined the efficacy of pCas4 vaccination in limiting disease burden during chronic infection in C57BL/6 mice. We aimed to measure bacterial burden as well as lung pathology and the immune cell composition of the lungs in mice vaccinated with pCas4 at 12 weeks post infection (Fig. 2A). Despite the enhanced T cell responses observed in pCas4 vaccinated mice at 4 weeks post-infection (Fig. 1D), we did not observe a reduction in bacterial burdens in either the lungs or spleens in vaccinated mice at 12 weeks post-infection (Fig. 2B). However, the gross pathology of lungs from pCas4-vaccinated mice appeared less than that in mock-vaccinated animals (data not shown). H&E staining of lung sections from mock- and pCas4-vaccinated mice revealed that there were fewer lymphoid aggregates in pCas4-vaccinated mice (Fig. 2C). Quantification of these structures using 2-dimensional image analysis using QuPath confirmed this observation revealing both fewer and smaller lymphoid aggregates in pCas4-vaccinated animals (Fig. 2D-E) [^33^]. Lymphoid aggregates are observed in chronic Mtb infection in mice and are a common feature of some granulomas in humans and non-human primates infected with Mtb [^12,34–36^]. These lymphoid aggregates mostly consist of B cells, and immunohistochemistry with the B cell marker B220 revealed that these structures were in fact primarily composed of B cells (Fig. 2F) [^34^]. Consistent with a reduction in these structures, we also observed a significant reduction in the frequency of B cells, as well as T cells, among the lung cells from pCas4-vaccinated animals (Fig. 2G-H). Overall, we interpreted these data to suggest that pCas4 vaccination may be limiting the immunopathology in mice without affecting the bacterial burdens, consistent with the principle of infection tolerance.

**Figure 2.**
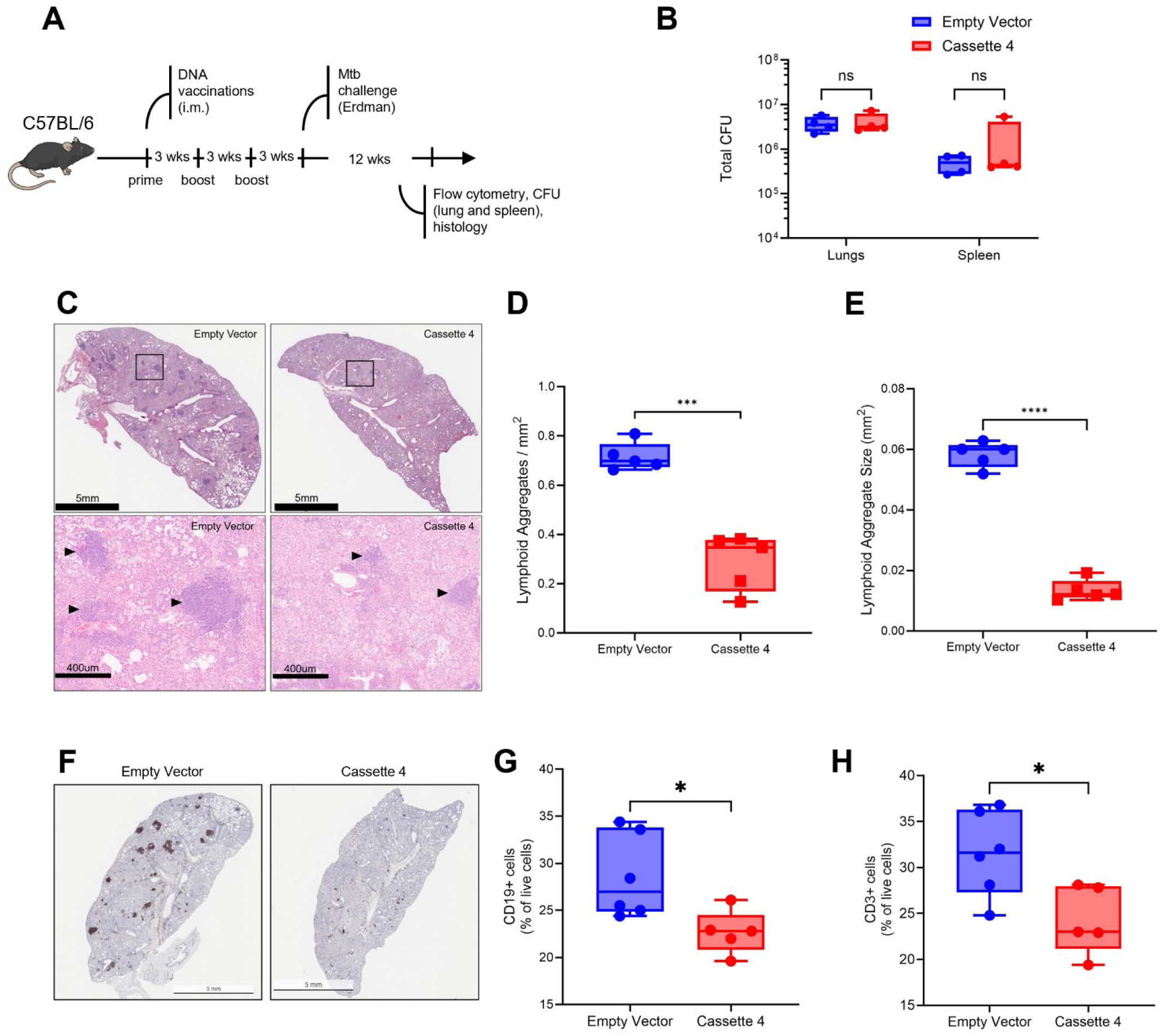
Impact of Cassette 4 vaccination on Mtb infection in C57BL/6 mice at 90 days post-infection. **A)** C57BL/6 mice were vaccinated with Cassette 4 and challenged with Mtb Erdman 3 weeks after the final vaccination. At 90 days post-infection, lungs were assayed for bacterial burden (CFU), immune cell composition (flow cytometry), and immune pathology (histology). **B)** No reduction in bacterial burden was observed in either the lungs or spleens of vaccinated mice. **C)** Histology revealed a reduction in the number and size of lymphoid aggregates in vaccinated animals. **D-E)** Quantification of lymphoid aggregates in H&E sections revealed a significant reduction in both frequency and size of lymphoid aggregates. **F)** Representative B220 staining reveals the lymphoid aggregates are mostly comprised of B cells. **E-G)** Flow cytometry revealed a significant reduction in the frequency of lymphocytes, CD19^+^ (B cells) and CD3^+^ (T cells), in vaccinated animals, which corresponds to the reduction in lymphoid aggregates observed. Data is representative of at least two independent experiments. Welch’s t tests were used to compare groups, and significant differences are indicated by a * (p-value < 0.05).

### Variable antigen vaccination reduces necrosis in SP140^-/-^ mice

With our observations of reduced immunopathology in C57BL/6 mice, we investigated the impact of pCas4-vaccination in hypersusceptible SP140^-/-^ mice [^29^]. These mice develop more severe and human-like pathology, with large necrotizing granulomas similar to those observed in humans [^12,29^]. Impacts to pathology or granuloma architecture without a change in bacterial burden have been observed in other mouse models [^28^]. We therefore hypothesized that pCas4-vaccination could reduce the overall pathology in SP140^-/-^ mice without impacting the enhanced bacterial replication observed in these mice. Since SP140^-/-^ mice are highly susceptible, we restricted our analyses to 5 weeks post-infection. Additionally, we employed intravenous injection of Sytox Green and whole lung imaging, as previously described, to more accurately quantify the extent of necrosis as a measure of lung pathology [^37^]. SP140^-/-^ mice were vaccinated with pCas4 or the pVAX1 empty vector and challenged with Mtb Erdman to assess the lung immune cell response, bacterial burdens in the lungs and spleens, and lung pathology (Fig. 3A). At 5 weeks post-infection we observed no significant difference in the bacterial burden in the lungs or spleens between mock- and pCas4-vaccinated mice (Fig. 3B). However, pCas4-vaccinated mice had reduced lung Sytox Green fluorescence, consistent with less necrosis (Fig. 3C). Quantification of whole lung fluorescence confirmed these observations showing a significant reduction in Sytox Green fluorescence in pCas4-vaccinated mice (Fig. 3E). H&E-stained sections confirmed the presence of large necrotizing lesions in mock-vaccinated mice, correlating with the high whole lung Sytox Green fluorescence (Fig. 3D). Mice vaccinated with pCas4 had large cellular lesions devoid of necrosis, but also lacking obvious organization (Fig. 3D).

**Figure 3.**
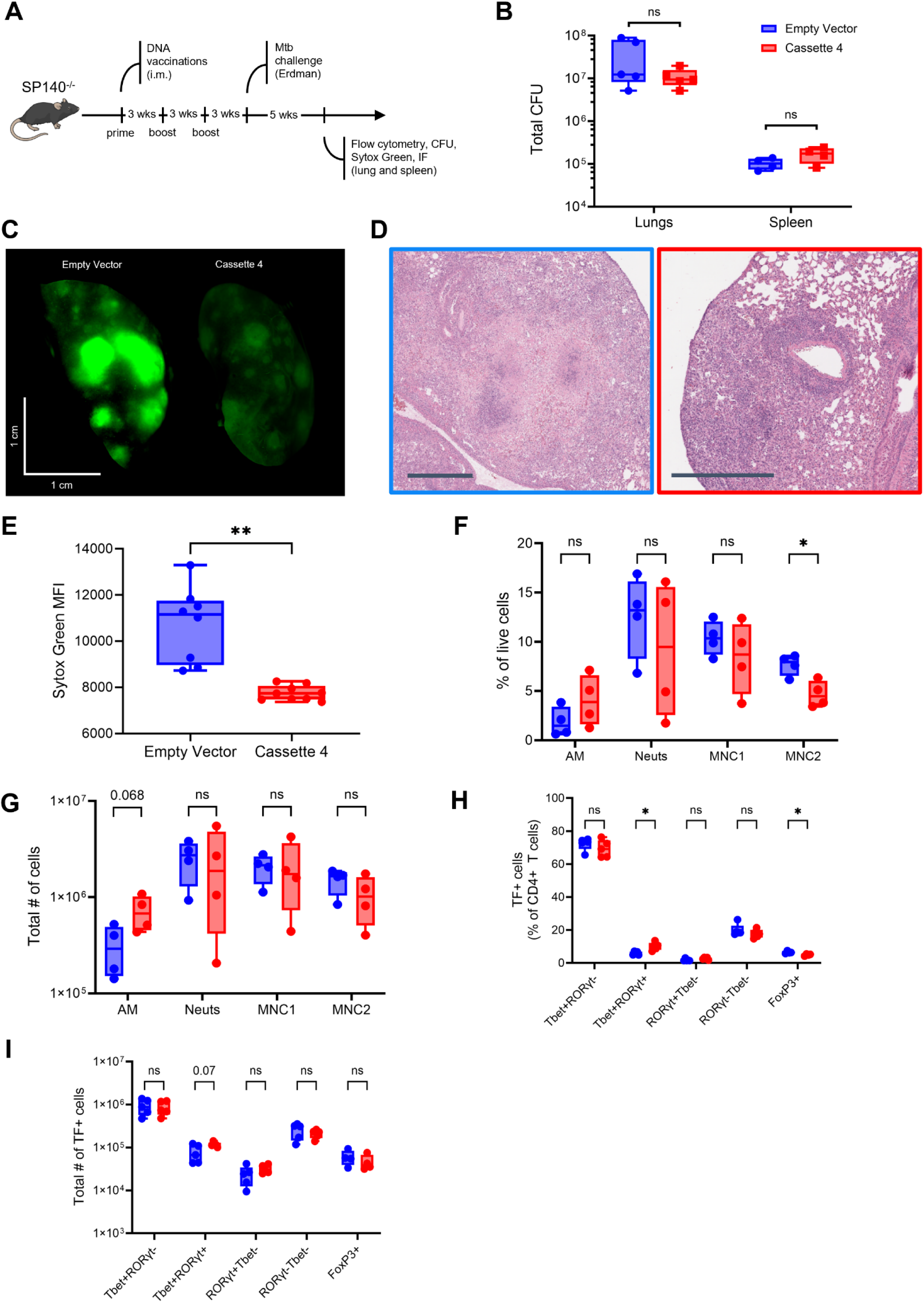
Cassette 4 vaccination reduces necrosis in susceptible SP140^-/-^ mice. **A)** SP140^-/-^ mice were vaccinated with Cassette 4 and challenged with Mtb Erdman. At 28 days post-challenge, mice were injected with Sytox Green 20 minutes prior to euthanasia to determine the extent of necrosis by whole lung imaging of the left lung. The right lungs were separated and used for quantifying bacterial burden (CFU) and immune cell composition (flow cytometry). **B)** Vaccinated animals did not have a significant reduction in CFU. **C)** Representative images of left lungs from Sytox Green-treated mice. **D)** Representative H&E sections of lesions from the same set of Sytox Green-treated lungs (scale bar = 250 µm). **E)** Quantification of whole lung Sytox Green MFI revealed a significant reduction in MFI in the lungs of vaccinated animals. **F)** Frequencies and **G)** absolute numbers of alveolar macrophages (AM), neutrophils (Neuts), and monocyte-derived cells (MNC1/MNC2) as a percent of total live cells. **H)** Frequencies and **I)** absolute number of CD4 T cell subsets expressing lineage defining transcription factors associated with Th1 and Th17 cells. Welch’s t tests were used to compare groups, and significant differences are indicated by a * (p-value < 0.05) or ** (p-value < 0.001).

The necrosis observed in SP140^-/-^ mice has been largely attributed to neutrophils [^29,38^]. Given the large differences in necrosis and lesion architecture in pCas4-vaccinated mice we predicted that pCas4-vaccination would reduce overall neutrophil numbers in the lungs. However, flow cytometry of lung cells revealed no significant impact of vaccination on frequency or total numbers of neutrophils in pCas4-vaccinated mice compared to mock-vaccinated mice (Fig. 3F-G). Instead, we observed a reduction in the frequency of MNC2 cells, which we previously demonstrated to be a more restrictive subset of monocyte-derived phagocytes during Mtb infection [^39^], and an increase in the number alveolar macrophages in pCas4-vaccinated mice relative to mock-vaccinated mice (Fig. 3F-G). With reduced lung pathology, it is likely that pCas4-vaccination prevents the destruction of healthy lung tissue and sustains higher numbers of alveolar macrophages while reducing the influx or differentiation of inflammatory monocyte-derived macrophages.

Since we identified an enhanced CD4 T cell response to the antigens in Cas4 after vaccination and challenge, we determined whether the lung CD4 T cell population was affected by pCas4-vaccination in SP140^-/-^ mice. Using flow cytometry and staining for lineage defining transcription factors, we determined the frequency and total numbers of Th1 (Tbet^+^), Th1/17 (Tbet^+^RORγt^+^), Th17 (RORγt^+^Tbet^-^), and Treg (FoxP3^+^) cells [^40^]. Mice that were vaccinated with pCas4 had a reduced frequency of Treg cells, possibly secondary to the reduced tissue damage observed by histology and whole lung Sytox Green fluorescence (Fig. 3H). Additionally, pCas4 vaccination induced a higher frequency of Tbet^+^RORγt^+^ (Th1/17) cells and a trend towards an increase in the total number of Tbet^+^RORγt^+^ cells, which have previously been associated with vaccine-mediated protection in mice (Fig. 3H-I) [^32^]. Although we identified an IL-17A response in the draining lymph nodes of C57BL/6 mice when stimulated with the Cas4 antigen, RimJ, we did not detect antigen-specific cells making IL-17A in the lungs (Fig. S3). This may be due to a technical limitation of our assay or indicate these cells are not making IL-17A despite expressing RORγt. Overall, we determined that an increased frequency of Th1/17 cells and reduction in a subset of monocyte-derived cells was associated with a reduction in lung necrosis in SP140^-/-^ mice.

### Vaccination with RVMA alters the immune response to Mtb infection

To further investigate the impact of pCas4-vaccination on the immune response to Mtb infection, we performed immunofluorescence microscopy of lung sections from DNA vector-only (mock)- and pCas4-vaccinated SP140^-/-^ mice. Due to the observation that pCas4-vaccinated mice had non-necrotic lesions, we aimed to assess the immune cell composition, as well as structural and bacterial composition, of the lesions. In SP140^-/-^ mice, the lesions that form during Mtb infection resemble human granulomas that consist of a necrotic neutrophil- and macrophage-rich core containing bacteria surrounded by a lymphocytic cuff [^12,38^]. We used antibodies against CD68, CD4, and Ly6G to identify macrophages, CD4 T cells, and neutrophils, respectively, within the lung lesions and divided each lesion into a core and cuff for analysis. Lesions from mock-vaccinated mice exhibited the expected composition of a large central necrotic core that stained positive for neutrophil and macrophage markers, with no intact cells and mostly acellular debris (Fig. 4A). Macrophages were present surrounding the necrotic core, along with CD4 T cells, which were almost entirely absent from the core of the lesion (Fig. 4A). In mice that were vaccinated with pCas4, the lesions contained swarms of intact neutrophils and pockets of macrophages, with CD4 T cells evenly distributed throughout the lesions (Fig. 4B).

**Figure 4.**
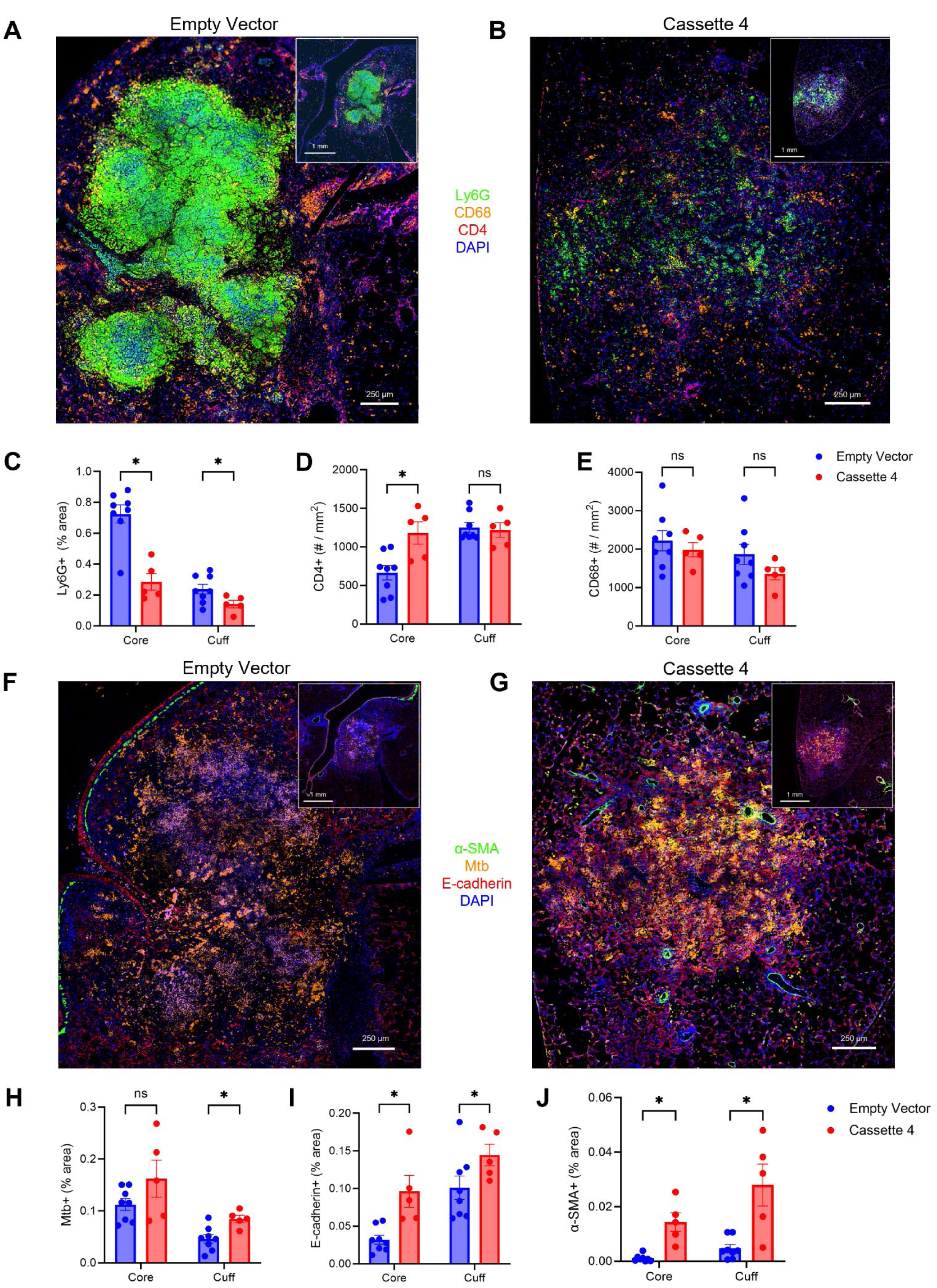
Lesion architecture in infected SP140^-/-^ mouse lungs are altered by Cassette 4 vaccination. Lung sections from vaccinated and mock-vaccinated SP140^-/-^ mouse lungs infected with Mtb Erdman 28 days before were stained with antibodies for **A-B)** neutrophils (Ly6G, green), macrophages (CD68, orange), and CD4 T cells (CD4, red), and counterstained with DAPI. Lesions were identified and subdivided into the core (inner 80%) and cuff (outer 20%). The **C)** percent area of neutrophil positive staining, **D)** CD4 T cell frequencies, and **E)** macrophage frequencies within each region were quantified. **F-G)** Structural components and bacteria within consecutive sections was also identified by staining for E-cadherin (red), myofibroblasts (α-SMA, green), and Mtb (ab905, orange), and counterstained with DAPI. The percent area within the core and cuff regions of the lesions with positive staining for **H)** Mtb, **I)** E-cadherin, **J)** and α-SMA was quantified. At least 5 lesions from at least 3 mock- and pCas4-vaccinated mouse lungs were imaged and used for analysis. Representative images contain inset of a larger 4X objective stitched view of the lung area around the lesion in the main image, which is a stitched image using the 20X objective. Welch’s t tests were used to compare groups, and significant differences are indicated by a * (p-value < 0.05).

Quantification of immune cell subsets across lesions from multiple mice revealed a significant reduction in neutrophil staining in pCas4-vaccination mice (Fig. 4C). Additionally, we determined that there was a significant increase in the number of CD4 T cells within the core of lesions, consistent with our observation that CD4 T cells were distributed throughout the non-necrotic lesions but absent from the core of necrotic lesions (Fig. 4D). No significant differences in the number of macrophages were identified (Fig. 4E).

Recent descriptions of granulomas from the susceptible C3HeB/FeJ mice have revealed that prior Mtb immunity reduced neutrophil-driven pathology and destruction of alveolar epithelium [^41^]. Mice vaccinated with pCas4 had lesions containing fewer neutrophils within the central core, and thus we hypothesized that these lesions would also exhibit less lung parenchymal tissue destruction. To identify lung tissue cells we stained tissue sections with antibodies against E-cadherin to mark alveolar epithelium and α-smooth muscle actin (α-SMA) to identify fibroblasts, as they have been observed in non-necrotic granulomas in other models [^42,43^].

Lesions from mock-vaccinated mice exhibited a loss of E-cadherin within the central core, which was expected given the acellular, neutrophil-rich appearance (Fig. 4F). Additionally, no obvious fibrosis was observed with little to no α-SMA staining observed (Fig. 4F). In pCas4-vaccinated mice, E-cadherin was observed throughout the lesions and not lost within the core (Fig. 4G). Interestingly, α-SMA staining resembling myofibroblasts associated with lung fibrosis was observed within the core and cuff of lesions from pCas4-vaccinated mice (Fig. 4G, Fig. S4C) [^42^]. Quantification confirmed the observed differences of significant increases in E-cadherin and α-SMA within the core of lesions from pCas4-vaccinated compared with mock-vaccinated mice (Fig. 4I-J). Consistent with the lack of difference in bacterial burdens between these two groups, we did not observe a difference in Mtb staining within the core of lesions between mock- and pCas4-vaccinated mice (Fig. 4H). Surprisingly, we observed a significant increase in Mtb staining within the cuff of lesions from pCas4-vaccinated mice (Fig. 4H). This may represent a poorer containment of the infection, normally contained mostly within the necrotic core, or a less organized lesion with less defined core and cuff regions. Regardless, these data demonstrate that pCas4-vaccination impacts TB lesion structures, in the absence of a significant reduction in bacterial burdens.

To further understand the role of pCas4 in mediating changes to pathology in SP140^-/-^ mice, we evaluated the immune cell composition in the lungs and the bacterial burden at an earlier time point. As we had only evaluated the impacts of pCas4 vaccination at 5-weeks post-infection, we determined whether differences in bacterial burden preceded changes in pathology and determined if differences in immune responses could be detected prior to development of severe lung necrosis in unvaccinated SP140^-/-^ mice (Fig. 5A). Consistent with our observations at later time points, we observed no differences in bacterial burdens in either the lungs or spleens between mock- and pCas4-vaccinated mice at this earlier time point (Fig. 5B). However, despite a lack of difference in bacterial burdens there were significant differences in the parenchymal immune cell composition in the lungs of pCas4-vaccinated mice. The changes in CD4 T cell subsets observed at 35 days post-infection were more apparent at 21 days post-infection. Specifically, we found a large increase in the frequency of RORγt-expressing CD4 T cells, both Tbet^+^ (Th1/17) and Tbet- (Th17) (Fig. 5C-D). This was accompanied by a decrease in the frequency of Tbet^+^ (Th1) CD4 T cells (Fig. 5C-D). The total number of these CD4 T cells mirrored the frequencies, without differences in the total number of CD4 T cells between the two groups (Fig. 5G).

**Figure 5.**
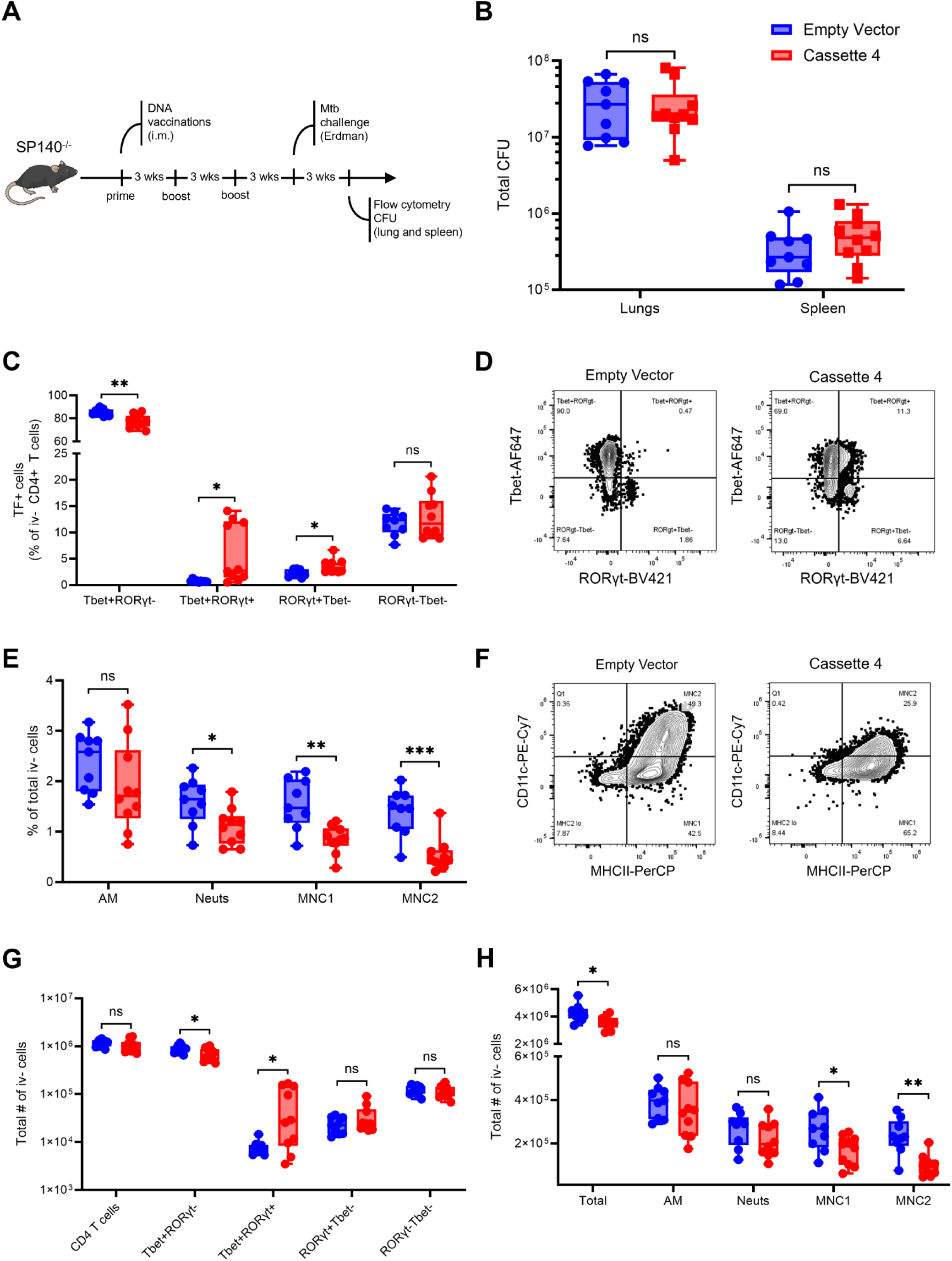
Cellular responses at 21 days post-infection differ in Cassette 4-vaccinated SP140^-/-^ mice. The immune cells in the lungs of mice vaccinated with Cas4 or the empty vector were assessed at 21 days post-infection. **A)** Experimental timeline of day 21 immune cell phenotyping. **B)** Bacterial burden assessed by CFU in the lungs and spleens of mock and vaccinated animals. **C)** Frequency of CD4 T cell subsets based on the expression of lineage-defining transcription factors, RORγt and Tbet, as a percent of iv^-^ CD4^+^ T cells. **D)** Representative gating of transcription factor positive CD4 T cells. **E)** Frequencies of alveolar macrophages (AM), neutrophils (neuts), mononuclear cell subset 1 (MNC1) and mononuclear cell subset 2 (MNC2) as a percent of total iv-cells. **F-G)** Approximate total cell numbers for CD4 T cell subsets and myeloid cell subsets including total CD4^+^ T cells and total lung cells. At least 5 mice per group were analyzed. Multiple Welch’s t tests with multiple testing correction when required was used to compare groups, and significant differences are indicated by * (p-value < 0.05), ** (p-value < 0.001), or *** (p-value < 0.0001).

In addition to the changes in CD4 T cells, we observed a significant reduction in the frequency of neutrophils and both monocyte-derived macrophage subsets, MNC1 and MNC2, in pCas4-vaccinated mice (Fig. 5E). The monocyte-derived cells in pCas4-vaccinated mice also had a reduction in the expression of CD11c (Fig. 5F), which may represent a less inflammatory cell state [^44,45^]. The total number of neutrophils was not significantly different in pCas4-vaccinated mice, but there was a significant reduction in the total number of both monocyte-derived cell subsets in pCas4-vaccinated mice corresponding to a small decrease in the total number of cells isolated from the lungs (Fig. 5H). These changes may be leading to the reduction in lung necrosis in pCas4-vaccinated mice. The reduction in neutrophil frequency observed at 21 days post-infection is likely not observed later due to the neutrophil cell death in mock-vaccinated mice reducing the number of neutrophils that are able to be detected by flow cytometry of live cells from digested lung tissue.

### B cells are dispensable for pCas4-mediated protection

B lymphcytes have been shown to be protective in Mtb infection in mice and macaques [^46^]. Therefore, we sought to determine if the effects of pCas4 vaccination on necrosis in SP140^-/-^ mice are mediated by B cells. To eliminate B cell responses to vaccination without affecting CD4 T cell responses, we used an anti-CD20 antibody to deplete B cells during vaccination (Fig. 6A). By depleting B cells during the vaccination regimen, we aimed to eliminate the induction of antigen specific B cells, while allowing B cell numbers to recover to baseline prior to Mtb challenge. At 21 days post-infection, B cell numbers had mostly recovered with only a trend towards reduced parenchymal B cells in anti-CD20 antibody-treated mice (Fig. S5). Depletion of B cells during vaccination did not impact the reduction in necrosis as measured by Sytox Green fluorescence in pCas4-vaccinated mice (Fig. 6B-C). Additionally, no difference in the lung bacterial burdens was observed at either 21- or 28-days post-infection (Fig. 6D). A reduction in the frequency of Tbet^+^ CD4 T cells was observed in pCas4-vaccinated B cell depleted mice similar to untreated pCas4-vaccinated mice (Fig. 6E-F). Similar reductions in monocyte-derived cell frequencies were also observed in both the pCas4-vaccinated anti-CD20 antibody treated mice and pCas4-vaccinated control mice (Fig. 6G-H). These data suggest B cells are dispensable for pCas4-mediated protection.

**Figure 6.**
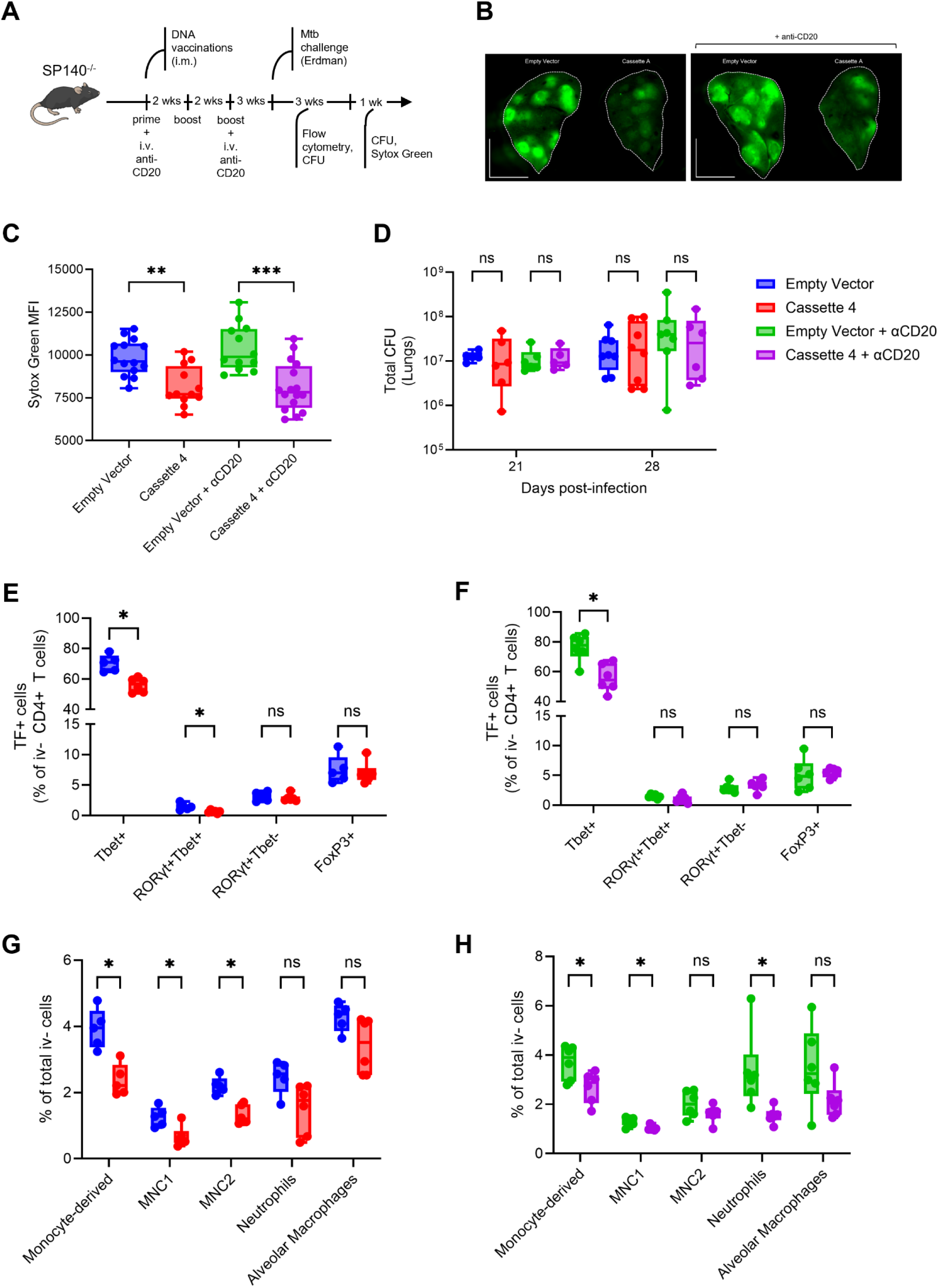
Vaccine-induced B cell responses are not required for Cassette 4-mediated reduction in necrosis in SP140^-/-^ mice. SP140^-/-^ mice were injected with anti-CD20 antibody in conjunction with vaccination with empty vector (mock) or Cas4 to prevent the generation of antigen-specific B cells. **A)** Experimental outline showing vaccine and anti-CD20 treatment. Mice were euthanized at both 21- and 28-days post-infection for lung immune cell analysis and Sytox Green injection for determining lung necrosis, respectively. **B)** Representative Sytox Green lung images and **C)** quantification of Sytox Green fluorescence. **D)** Bacterial burden in the lungs of mice quantified by CFU. Frequencies of CD4 T cell subsets based on lineage-defining transcription factor expression as a percent of total iv-CD4^+^ T cells for **E)** B cell replete and **F)** B cell depleted mice. Frequencies of myeloid cell subsets as a percent of total iv-cells for **G)** B cell replete and **H)** B cell depleted mice. At least 5 mice per group per timepoint were analyzed. Multiple Welch’s t tests with multiple testing correction when required was used to compare groups, and significant differences are indicated by * (p-value < 0.05), ** (p-value < 0.001), or *** (p-value < 0.0001).

### A vaccine encoding for RimJ alone is sufficient to reduce necrosis in SP140^-/-^ mice

Vaccination with pCas4 induced immune responses to 3 of the 4 RVMA, but RimJ was of special interest due to the amino acid substitution between H37Rv and Erdman Mtb strains. To test whether vaccination with RimJ alone could recapitulate the effects observed after pCas4 vaccination, we vaccinated SP140^-/-^ mice with the pVAX1-RimJ plasmid and challenged with Mtb Erdman (Fig. 7A). RimJ vaccination significantly reduced Sytox Green staining in whole lungs of SP140^-/-^ mice compared with mock-vaccination (Fig. 7B-C). No significant change in the bacterial burden in lungs or spleens of mock- or RimJ-vaccinated mice was observed, consistent with the observations made using pCas4 (Fig. 7D). The immune cell composition was only measured at 28 days post-infection, and most differences were more subtle than what was observed for pCas4-vaccination at 21 days post-infection. Nonetheless, RimJ-vaccinated mice had significantly increased frequencies and numbers of RORγt^+^ CD4 T cells, as was observed in pCas4-vaccinated mice (Fig. 7E-F). However, no significant reduction in monocyte-derived cells was observed at this time-point in RimJ-vaccinated animals (Fig. 7G-H). It is important to note that this assessment did not include discrimination between intravascular and extravascular cells, and thus the cell populations analyzed were not restricted to parenchymal cells alone, which may have masked subtle differences. Although we observed a significant increase in natural killer (NK) cells in RimJ-vaccinated mice, it is unclear how vaccination with RimJ might impact the recruitment of NK cells or what role NK cells might have in limiting the pathology. NK cells have been shown to recognize Mtb and may play a role in helping to limit pathology at the this time point [^47^]. Overall, RimJ-vaccination is able to recapitulate most aspects of pCas4-vaccination in SP140^-/-^ mice and thus represents an antigen for further investigation of the differences in T cell responses to this class of antigens and how naturally occurring T cell epitope-disrupting substitutions impact vaccine efficacy.

**Figure 7.**
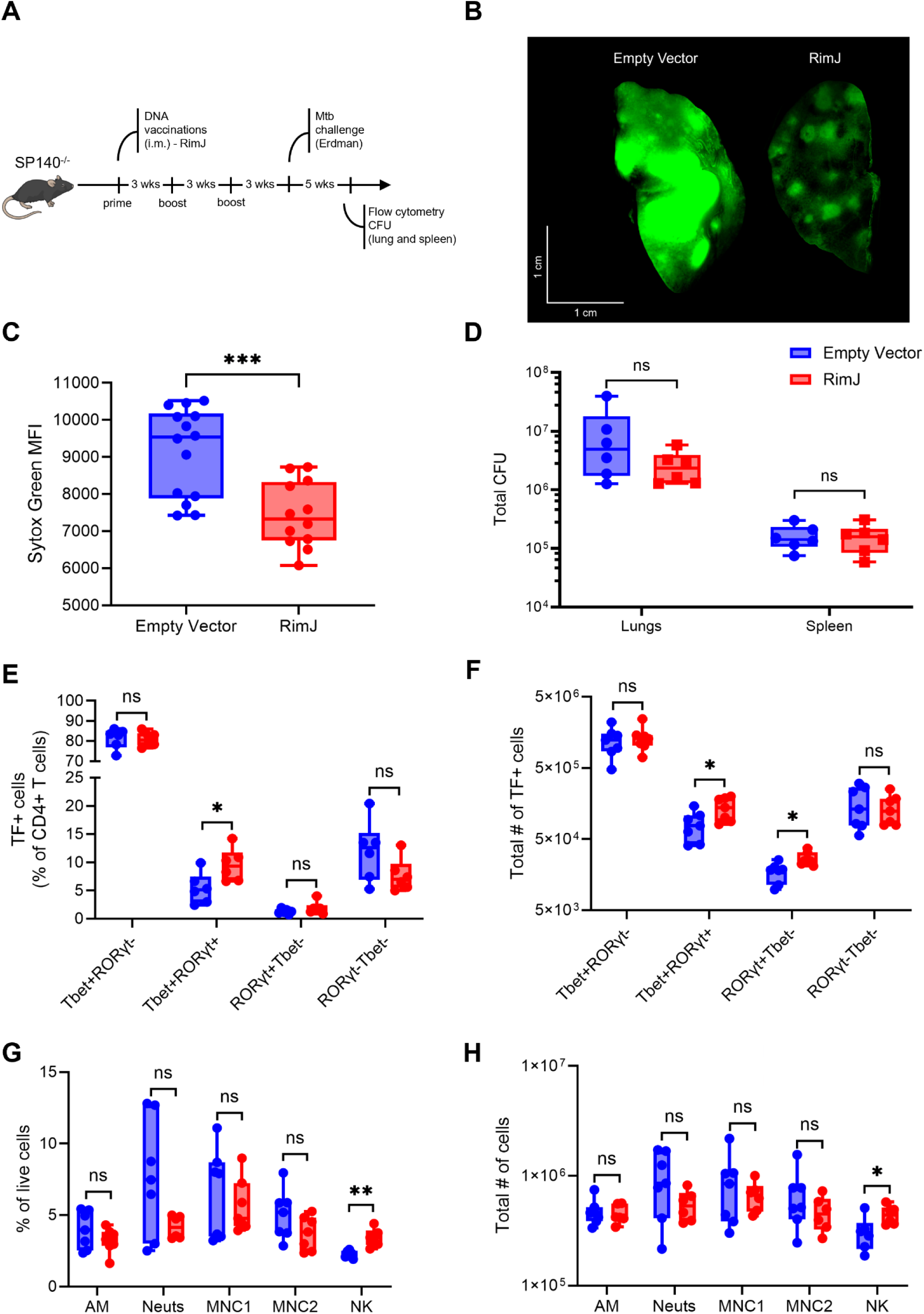
A DNA vaccine encoding only RimJ is sufficient to reduce necrosis in SP140^-/-^ mice. A DNA vaccine consisting of only RimJ, from Mtb Erdman, was used to vaccinate SP140^-/-^ mice. **A)** Experimental timeline for vaccination and challenge of SP140^-/-^ mice with RimJ and matched Mtb Erdman. **B)** Representative Sytox Green lungs from mock- and RimJ-vaccinated mice at 35 days post-infection and **C)** quantification of Sytox Green fluorescence. **D)** Bacterial burden in the lungs and spleens of mock- and RimJ-vaccinated mice at 35 days post-infection determined by CFU. **E)** Frequencies of CD4 T cells expressing the lineage-defining transcription factors Tbet and RORγt as a percent of total CD4 T cells, and **F)** absolute numbers of CD4 T cell subsets in the lungs. **G)** Frequencies of non-CD4 T cell subsets, which includes natural killer (NK) cells, as a percent of total iv-cells. **H)** Total numbers of specified cell subsets in the lungs. At least 5 mice per group were analyzed. Multiple Welch’s t tests with multiple testing correction when required was used to compare groups, and significant differences are indicated by * (p-value < 0.05), ** (p-value < 0.001), or *** (p-value < 0.0001).

To further characterize the CD4 T cell response to RimJ, we transferred CD4 T cells from vaccinated congenic mice to allow for phenotyping and tissue localization in infected SP140^-/-^ mice. Spleen cells from vaccinated CD45.1 congenic mice were stimulated with peptides and selected by magnetic enrichment for antigen-responsive CD154^+^ cells. CD154^+^ cells were then transferred into recipient CD45.2 SP140^-/-^ mice one day prior to aerosol infection with Mtb Erdman expressing the fluorescent protein zsGreen. We then assessed the presence, phenotype, and localization of CD45.1^+^ CD4^+^ T cells 28 days post-infection using flow cytometry and immunofluorescence (Fig. 8A and E). We identified CD45.1^+^CD4^+^ T cells in the lungs by both modalities (Fig. 8A and E). Transcription factor staining revealed a significant reduction in the frequency of Tbet^+^ cells among the CD45.1^+^ CD4 T cells compared with endogenous (CD45.1^-^) CD4 T cells (Fig. 8B-C). Notably, RORγt^+^ cells were also significantly reduced among transferred cells as compared to endogenous CD4 T cells. (Fig. 8B-C). We confirmed that CD45.1^+^ CD4 T cells were RimJ-specific by restimulation with RimJ peptides and intracellular cytokine staining for IFNγ, which revealed responses among the CD45.1^+^ CD4 T cells only to RimJ peptides and not to ESAT-6 peptides (Fig. 8D). Additionally, staining for IL-10 revealed no detectable IL-10 expression among the transferred cells, which indicates that RimJ-specific CD4 T cells are not a significant source of this immunosuppressive cytokine (Fig. 8D). Immunofluorescence of lung tissue sections revealed large non-necrotic lesions containing CD45.1^+^CD4^+^ cells within the lesions near Mtb^+^ cells (Fig. 8E). These data indicate that vaccine-induced RimJ-specific CD4 T cells respond and traffic to the lungs during Mtb infection and likely exhibit a unique phenotype in the context of the bulk CD4 T cell response to Mtb infection.

**Figure 8.**
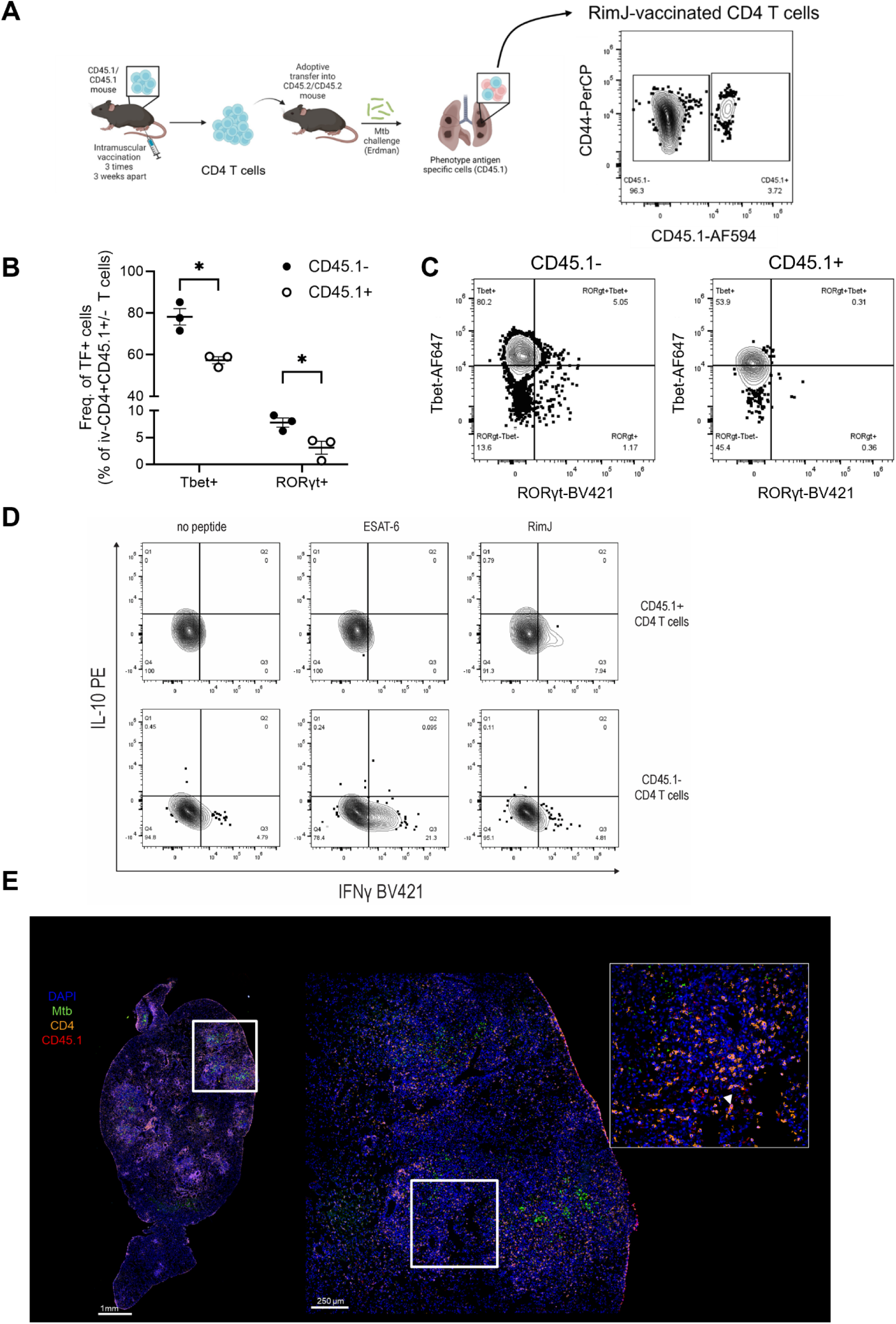
Vaccine-specific CD4 T cells from RimJ-vaccinated mice are present in the lungs during Mtb infection. **A)** Congenic CD45.1 mice were vaccinated with a DNA vaccine encoding for RimJ from Mtb Erdman. Vaccine-specific CD4 T cells were isolated from spleens by stimulation with RimJ peptides and subsequent enrichment for CD154^+^ cells by magnetic selection. 5×10^5^ CD154^+^ cells were transferred into CD45.2 SP140^-/-^ mice the day before aerosol infection with Mtb Erdman expressing the fluorescent protein zsGreen. At 28 days post-infection, lungs were assessed by flow cytometry and immunofluorescence to determine CD45.1^+^CD4^+^ vaccine CD4 T cell presence and phenotype. **B)** Transcription factor staining revealed a significant reduction in the frequency of Tbet^+^ and RORγt^+^ cells among the CD45.1^+^ CD4 T cells as compared to the endogenous (CD45.1^-^) CD4 T cells. **C)** Representative flow cytometry gating of transferred (CD45.1^+^) and endogenous (CD45.1^-^) CD4 T cells. **D)** Peptide stimulation and intracellular cytokine staining revealed a specificity for RimJ peptides by IFNγ^+^ cells and low levels of background IFNγ production and no IL-10 expression. **E)** Immunofluorescence of lung tissue from a mouse that received vaccine-specific CD4 T cells revealed the presence of CD4^+^ (orange) CD45.1^+^ (red) cells within Mtb-containing (green) lesions (white arrows in inset) and characteristic large non-necrotic lesions. Experimental layout was created using Biorender. Welch’s t tests were used to compare frequencies between endogenous and transferred T cells, and significant differences are indicated by a * (p-value < 0.05).

## Discussion

We generated DNA vaccines encoding RVMA, which were previously identified by their presence of nonsynonymous mutations within human T cell epitopes suggestive of immunological selection pressure [^16,17^] and found that vaccination with DNA encoding one or more RVMA prior to Mtb infection limits immune pathology without reducing the bacterial burdens. Vaccination with a DNA vaccine encoding for a fusion of 4 variable antigens reduced lymphoid aggregate formation in resistant C57BL/6 mice and limited formation of necrotic lesions in susceptible SP140^-/-^ mice. Additionally, these non-necrotic lesions in pCas4-vaccinated mice had reduced lung tissue destruction and signs of myofibroblast presence. Corresponding reductions in neutrophils and increased CD4 T cell penetration into the core of lesions in pCas4-vaccinated mice were consistent with other studies in susceptible mice [^41^]. Early differences in immune responses precede these changes in pCas4-vaccinated mice. Reductions in neutrophils as well as monocyte-derived cells, coupled with changes to CD4 T cell subsets, may all contribute to preventing formation of necrotic lesions. B cells appear to be dispensable in the context of vaccination with pCas4. Further, vaccination with a DNA vaccine encoding for RimJ alone is sufficient to promote this reduction in necrosis and changes to the CD4 T cell subsets responding to the infection.

Aside from a higher rate of nonsynonymous mutations within T cell epitope regions of the genes, it is unclear what might be different about these variable antigens as compared to more common vaccine antigens, such as ESAT-6. The most obvious difference is the lack of secretion for these antigens as compared to the classical immunodominant antigens. This may be the reason we observed a more modest T cell response to RVMA in response to Mtb infection of unvaccinated mice compared to the immunodominant antigens, ESAT-6 and Ag85B. While other work has shown that vaccination with ‘subdominant’ antigens can induce protection, it is unclear how being subdominant impacts T cell responses to RVMA [^31^]. Targeting T cell responses specifically to subdominant epitopes within ESAT-6 in mice was able to improve quality of the CD4 T cell response to Mtb infection [^48^]. Thus, these antigens inducing subdominant CD4 T cell responses may generate CD4 T cell responses that reduce lung pathology while not increasing control of bacterial replication.

pCas4- or RimJ vaccination increased RORγt^+^ CD4 T cells. RORγt^+^ CD4 T cells are increased in frequency in people who resist TB infection and in vaccinated mice with enhanced restriction of Mtb, and thus may contribute to protective CD4 T cell responses [^32,49^]. While observed increases in the frequency of RORγt^+^ CD4 T cells in the lungs of pCas4- and RimJ-vaccinated mice, we did not able detect IL-17A expression by ICS in the lungs of infected mice after stimulation with RimJ peptides (Fig. S3B). However, we did detect IL-17A and IFNγ production in response to RimJ-stimulation in the draining lymph node at early time points post-infection (Fig. S3A). Thus, CD4 T cell responses to RimJ may represent a natural ex-Th17 response that may be impacting Mtb-induced tissue damage without restricting replication. Indeed, we can detect ex-Th17 cells in the lungs of infected mice using an *Il17a-cre;*Ai14 reporter mouse to fate-map cells that express IL-17A (Fig. S3C-D), and these cells have been implicated in lung bacterial clearance [^50,51^]. Further studies will be needed to elucidate the impact of ex-Th17 cells in the context of Mtb infection.

A consequence of choosing sequence-variable antigens for these experiments was identifying a naturally occurring substitution in RimJ in the Mtb Erdman strain that was predicted to impact MHCII binding for H2-IA^b^. We found that the Erdman RimJ peptide variant elicited lower frequencies of CD4 T cells in H37Rv-infected mice and allowed for recognition of a secondary epitope in mice vaccinated with the Erdman variant (Fig. S2). RimJ vaccination alone was able to recapitulate the effects of pCas4-vaccination, and thus responses to RimJ likely account for many of the effects mediated by pCas4-vaccination. RimJ can serve as a model for future studies that aim to investigate the impact of antigenic variation on the CD4 T cell response to Mtb and vaccine efficacy.

Immune pathology associated with Mtb infection in mice may not be linked to bacterial burden in all contexts. Indeed, it has been observed in some mouse models, such as mice deficient in the intracellular adhesion molecule 1 (ICAM-1) gene, that changes in pathology can be independent of bacterial burden as described here [^52,53^]. It is even possible to achieve increased pathology with lower bacterial burdens in the context of IFNγ-overexpression [^27^]. Identifying a context in which vaccination with specific antigens can create a similar decoupling of pathology and bacterial replication has important consequences for preclinical testing of future Mtb vaccines. Ensuring that bacterial burden is not used as a sole method of ruling out a vaccine may be important for identifying vaccines that achieves substantial efficacy in humans. Additionally, the fact that CD4 T cell responses to Mtb may promote pathology and possibly transmission warrants caution and consideration in early-stage trials. Further studies are needed to determine whether evolutionary pressures are selecting for Mtb that induces strong CD4 T cell responses to promote transmission, and whether RVMA induce CD4 T cell responses that protect the host from pathology and enhanced transmission.

## Materials and Methods

### Plasmids

Vaccine cassettes were made by restriction cloning synthetic gene fragments (Genewiz) into the pVAX1 vector backbone (Invitrogen). Sequences were derived from Mtb Erdman strain and mammalian codon optimized. Four variable antigens Rv0990c, Rv0010c, RimJ (Rv0995), and Rv2719c were used to construct the vaccine vector used in these studies, named Cassette 4 (Cas4). NEB 5-alpha (NEB) *E. coli* was transformed with resulting vaccine plasmids and propagated in LB broth containing 50 µg/ml kanamycin. Vaccine plasmid for immunizations was prepared from 2L of overnight culture by Gigaprep (Zymo Research) with subsequent endotoxin removal. Vaccine plasmid diluted to a concentration of 2 μg/μl in sterile phosphate buffered saline (PBS) and stored at −20°C.

### Mice

C57BL/6 mice were obtained from Jackson Laboratory. SP140^-/-^ mice were obtained from the laboratory of Russel Vance and bred under specific-pathogen-free conditions at the University of California, San Francisco [^29^]. Congenic CD45.1 mice (B6.SJL-Ptprca Pepcb/BoyJ, strain #:002014) were obtained from Jackson Laboratory. Mice aged 8-12 weeks for used for experiments, and mice infected with Mtb were housed in the Animal Biosafety Level 3 facility. All animal protocols used were approved by the University of California, San Francisco, Animal Care and Use Committee.

### Mycobacterial strains, culture, and aerosol infections

Mtb Erdman, obtained from ATCC (ATCC 35801), Mtb H37Rv, and Mtb H37Rv expressing zsGreen, generated previously by our lab, were propagated in Middlebrook 7H9 (BD) supplemented with 10% (v/v) ADC (albumin, dextrose, catalase), 0.05% Tween-80, and 0.2% glycerol [^39^]. For aerosol infections, a frozen stock of mid-log phase bacteria was thawed and passed through a 25-G needle to ensure a single cell suspension. Bacterial stocks were diluted to an approximate concentration in water to yield an infection with 50-100 CFU per mouse. Mice were infected with Mtb using an inhalation exposure unit from Glas-Col as described previously [^39^]. Initial infectious dose was confirmed by homogenizing lungs from mice in PBS + 0.05% Tween-80 immediately after the aerosol procedure. Homogenates were plated on 7H11 agar plates containing PANTA (BD) to enumerate CFU [^54^].

### Immunizations and antigen stimulations

Mice were immunized in the right tibialis anterior muscle as described previously [^55–59^]. Briefly, 50 µl of vaccine plasmid (100 μg total plasmid) was injected using a 28G ½” 0.5 ml insulin syringe. Mice were immunized three times, three weeks apart. All experiments were performed three weeks after the final vaccination.

For assessing immunogenicity in vaccinated animals, mice were euthanized, and spleens were collected into RPMI-1640 containing 10% (v/v) heat-inactivated fetal bovine serum, 10mM HEPES, penicillin/streptomycin, and 50µM 2-mercaptoethanol (complete RPMI). Spleens were mashed through a 70µm strainer using the back of 3ml syringe to generate a single cell suspension. Red blood cells (RBCs) were lysed using ACK lysis buffer (Gibco). Spleen cells were cultured in complete RPMI at 37°C + 5% CO_2_ in 96-well round bottom plates with 10µg/ml pooled overlapping peptides (18-mers, overlapping by 11 amino acids) from Genscript or individual peptides where specified. For IFN-γ secretion, cells were incubated for 72 hours before supernatants were collected for analysis by IFN-γ ELISA (Invitrogen).

For flow cytometry, cells were incubated for 1 hour prior to the addition of eBioscience protein transport inhibitor cocktail (Invitrogen) and an additional 5-hour incubation. Stimulated cells were then stained with Live-or-Dye750-777 (Biotium) and anti-CD16/32 (clone 2.4G2; Fc block) for 15 minutes at 4°C. After washing, cells were stained for surface markers in FACS buffer (PBS with 2% HI-FBS (v/v) for 30 minutes at 4°C containing the following fluorochrome-conjugated antibodies: CD3 (17A2), CD4 (GK1.5), CD8 (53-6.7), CD19 (6D5), and CD44 (IM7). After surface staining, cells were fixed and permeabilized using Cytofix/Cytoperm (BD Biosciences) and stained in BD Perm / Wash buffer containing antibodies for IFNγ (XMG1.2) and TNF (MP6-XT22) for 30 minutes at 4°C. Samples were acquired using an Aurora spectral flow cytometer (Cytek Biosciences) and gated as shown in Supplemental Figure 1.

### Tissue harvests and flow cytometry

Where specified, mice were anesthetized by inhalation of isoflurane and retro-orbitally injected with 1 µg of anti-CD45 antibody (clone 30-F11) in 100 µl of PBS, 3 minutes before euthanasia with CO_2_ inhalation and cervical dislocation. Lungs were collected into Hanks’ Balanced Salt Solution (HBSS) containing 50 µg Liberase TM (Sigma) and 30 µg/ml DNase I (Sigma) and incubated at 37°C for 30 minutes, then processed with a gentleMACS dissociator (Miltenyi). To obtain a single cell suspension, the resulting homogenates were passed through a 70 µm strainer. Spleens were also collected into complete RPMI as described for single cell suspensions or PBS + 0.05% (v/v) Tween-80 for enumerating CFU. Aliquots of lung and spleen samples were taken prior to further processing for enumerating CFU by serial dilution and plating on 7H11 agar plates. Lung single cell suspensions were further processed by centrifugation and RBC lysis using ACK lysis buffer (Gibco).

For flow cytometry analysis, lung single cell suspensions were first stained with Live-or-Dye750-777 (Biotium) and Fc block for 15 minutes at 4°C. Lung cells were then stained in FACS buffer with two sets of antibodies for analyzing myeloid cell populations as described previously [^39^] and CD4 T cell subsets by lineage defining transcription factors. For myeloid cell analysis, cells were stained in FACS buffer containing a cocktail of fluorochrome-conjugated antibodies, as described previously [^39^], for 30 minutes at 4°C. For CD4 T cell analysis, cells were stained in FACS buffer for 30 minutes at 4°C containing the following fluorochrome-conjugated antibodies: CD3 (17A2), CD4 (GK1.5), CD8 (53-6.7), CD19 (6D5), and CD44 (IM7). For CD4 T cell subset analysis, cells were then fixed and permeabilized using the eBioscience FoxP3 / Transcription Factor Staining Buffer Set (Invitrogen) prior to being incubated with fluorochrome-conjugated antibodies targeting Tbet (4B10), RORγt (Q31-378), and FoxP3 (MF-14). All cells were analyzed on an Aurora Spectral flow cytometer (Cytek Biosciences). Myeloid cell subsets were identified as described previously [^39^], and CD4 T cell subsets were identified as live > single cells > CD3^+^CD19^-^ > CD4^+^CD8^-^ > CD44^+^ (Fig. S1).

### Lung tissue histopathology, cryosectioning, and immunofluorescence

Mice were euthanized, as described above, and lungs were harvested directly into 10% neutral buffered formalin for histopathology. After 48 hours, fixed lungs were shipped to Histowiz for processing. All H&E staining and immunohistochemistry was performed by Histowiz. Histopathology was assessed using QuPath to quantitate lymphoid aggregates in lungs from at least 5 mice per group [^33^]. Lymphoid aggregates were identified by training a pixel classifier to identify lymphoid-rich regions (dense lymphocyte aggregates) within the lung sections, and B220 IHC confirmed lymphocyte identification by H&E staining. The number of lymphoid aggregates in the lungs was used to calculate the frequency (number / total lung area), and the size of each identified lymphoid aggregate was also measured.

For cryosectioning and immunofluorescence, mice were euthanized, as described above, and lungs were inflated with PBS containing 50% (v/v) OCT compound through the trachea. Lungs were embedded in OCT and flash frozen in liquid nitrogen. Blocks were cut into 6 µm sections by cryostat (Leica Biosystems), and sections were transferred to CFSA 1X slides (Leica Biosystems) using adhesive tape windows (Leica Biosystems). Sections were crosslinked to the slides via UV exposure and fixed in acetone for 15 minutes. After drying, slides were stored at −20°C until staining.

For immunofluorescence staining, slides were rehydrated for 10 minutes in PBS before blocking with PBS + 0.5% BSA for 30 minutes at room temperature. Sections were then stained in PBS + 0.5% BSA containing fluorochrome-conjugated antibodies against CD4 (GK1.5), CD68 (FA-11), and Ly6G (1A8), or E-Cadherin (DECMA-1) and α-SMA (1A4) for 2 hours at room temperature. E-cadherin and α-SMA sections were also stained with a primary unconjugated antibody against Mtb (ab905, Abcam) and subsequently stained with a fluorochrome-conjugated isotype specific secondary antibody for 1 hour at room temperature. All sections were washed and stained with DAPI before coverslips were mounted with VECTASHIELD Vibrance Antifade Medium (Vector Laboratories). Images were captured using the 4X and 20X objective of a Nikon Ti inverted microscope with a DS-Qi2 camera. Individual images were stitched together to cover entire areas using NIS Elements software (Nikon).

Lesions were manually identified from whole lung images using the 4X objective and were further imaged using the 20X objective. At least 5 mice per group were imaged. Images were imported into QuPath for analysis [^33^]. Lesions identified by Mtb staining or Ly6G-rich regions were sub-divided into core and cuff region by manually annotating the central 80% of the lesion as the core and the outer 20% as the cuff (Fig. S4A-B). Analysis was performed by utilizing a pixel thresholder in QuPath to mask cells or regions that had positive staining. Frequencies were determined by calculating the total number of identified events and dividing by the total area of the core or cuff regions for CD4 and CD68 staining, as those markers identified discrete cells. For Ly6G, which identified large necrotic areas, the total area masked was used to calculate the relative area of positive staining (positive staining area / total area) due to a lack of discrete objects. Similar analysis was performed for E-cadherin, α-SMA, and Mtb (ab905) staining.

### Sytox Green injection and lung imaging

To determine the extent of necrosis in infected mice, we utilized Sytox Green dye injection as described previously [^37^]. Briefly, infected mice were anesthetized by isoflurane inhalation and retro-orbitally injected with 100ul of a 50 μM solution of Sytox Green dye (Invitrogen), a live-cell impermeable dye that stains DNA, 20 minutes prior to euthanasia as described above. Lungs were immediately collected into PBS to wash, before being fixed in 10% neutral buffered formalin for 48 hours at 4°C. Fixed lungs were imaged using a Leica M205 FCA stereomicroscope and images captured using a Leica K5 monochrome camera. Post acquisition images were processed using LASX software (Leica). Image analysis was performed using ImageJ to quantify Sytox Green MFI of the left lung lobes from at least 5 mice per group. For experiments where Sytox Green injection was performed, the right lung lobes were utilized for flow cytometry and enumeration of bacterial burden by CFU, as described above.

### B cell depletion

To isolate the role of the vaccine-induced T cell response in preventing necrosis in SP140^-/-^ mice, B cells were depleted by anti-CD20 mAb injection (clone MB20-11, Bio X Cell) as described previously [^60^]. Briefly, mice were injected intravenously with 10 μg of anti-CD20 antibody in 100 μl of PBS 1 day before the first vaccination with either the pVAX1 empty vector (mock) or Cas4, and again 4 weeks later in the middle of the vaccine regimen, to deplete B cells and prevent a vaccine-specific B cell response from occurring. Mice were challenged after the final vaccination, approximately 5 weeks after the final anti-CD20 antibody injection, when B cell numbers should have recovered. Experiment was performed simultaneously along with a separate group of mice that did not receive anti-CD20 antibody. Tissue harvests for bacterial burden enumeration, flow cytometry, and the Sytox Green assay for determining whole-lung necrosis were performed as described above. Statistical comparisons were made between mock- and pCas4-vaccinated groups within the control and B cell depleted mice.

### CD4 T cell adoptive transfers and cell sorting

Congenic B6 CD45.1 mice were vaccinated as described above with RimJ. After vaccination, RimJ-specific CD4 T cells were purified from spleen cells of vaccinated mice by RimJ peptide stimulation and magnetic enrichment using the CD154 enrichment and detection kit (PE, mouse) (Miltenyi Biotec) according to the manufacturer’s instructions. Approximately 5×10^5^ purified CD154^+^ cells from vaccinated mice were transferred via retro-orbital injection into recipient SP140-/- mice under isoflurane anesthesia. The day after transferring vaccine CD4 T cells, the mice were infected via aerosol infection with Mtb Erdman expressing zsGreen, as described above. At the indicated time points, lung cells were prepared for cell sorting by staining for T cell flow cytometry as described above with the addition of a fluorochrome-conjugated antibody for CD45.1 (A20). Recipient mice were used for lung sectioning and imaging or antigen stimulation and ICS, as described above.

### Data analysis

Flow cytometry data were processed and analyzed using SpectroFlo 3.3 (Cytek) and FlowJo 10.10.0. For ICS experiments, data is background subtracted where the frequencies in the unstimulated sample were subtracted from the frequencies in the paired stimulated samples. Absolute numbers of cells were calculated using the number of events and volume of sample acquired by the cytometer. GraphPad Prism 10.4.1 (GraphPad) was used for generating plots and performing statistical analysis.

## Acknowledgements

We thank Irina Debnath for early contributions to this work. We also acknowledge the excellent technical assistance provided by I-Chang Chang and Lucas Chen. Microscopy data for this study was acquired at the UCSF Innovation Core at the Weill Institute for Neurosciences and the UCSF Center for Advanced Light Microscopy. Flow cytometry data for this study was generated with assistance from the UCSF Division of Experimental Medicine Core Immunology Lab. This research was supported by the NIH grants F31 AI172360-01 (Z.H.) and U01 AI166309 (J.D.E.). The funders had no role in study design, data collection or interpretation, or the decision to submit the work for publication.

## Supplemental Material

**Supplemental Figure 1.**
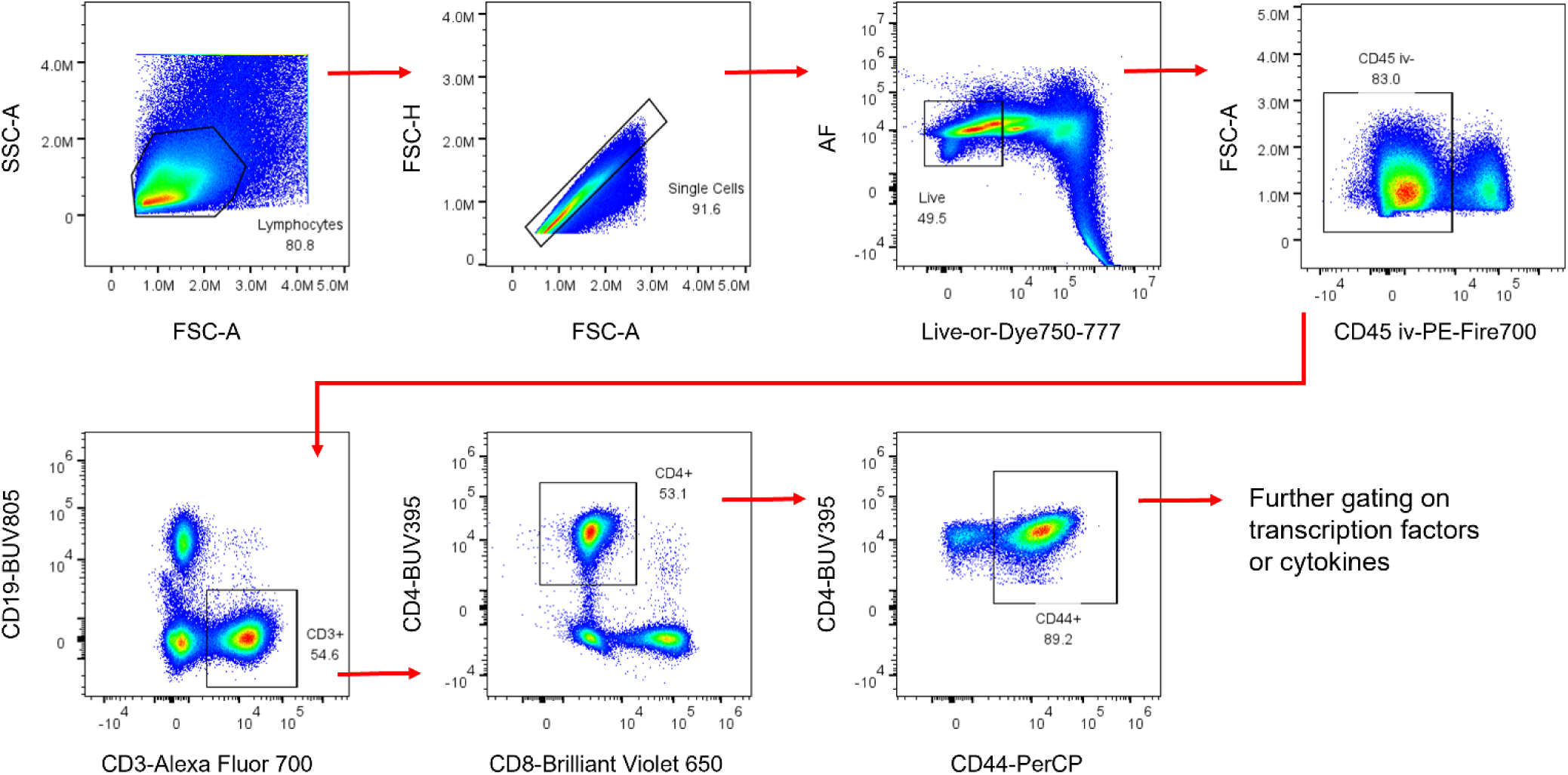
Gating strategy for assessing CD4 T cell subsets. For cytokine and transcription factor staining, only the CD4^+^ T cell CD44^+^ population was assessed.

**Supplemental Figure 2.**
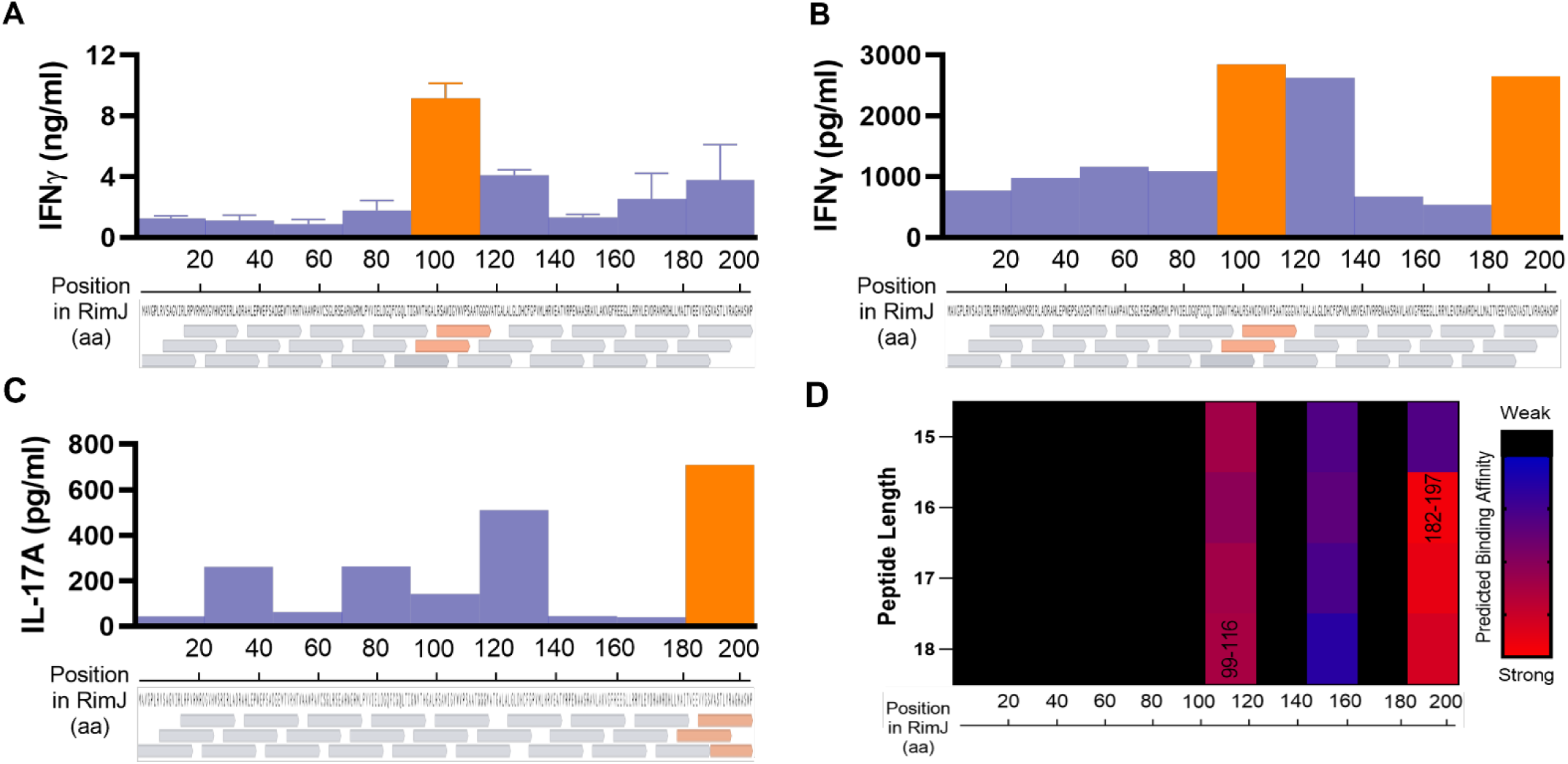
Epitope mapping for RimJ using pooled overlapping peptides in spleen and mediastinal lymph node. Pooled overlapping peptides (18-mers overlapping by 11) spanning the length of the RimJ antigen were used in small (2-4 peptides) pools at a concentration of 10 μg/ml to stimulate cells from spleen and lymph nodes from Mtb H37Rv-infected C57BL/6 mice *ex vivo* for 72 hours. A) Spleen cell responses as measured by IFNγ concentration in the cell culture supernatant revealed a dominant response to two peptides spanning amino acids 92-116. Data is mean±SEM (n = 3 mice). B-C) Mediastinal lymph nodes from 5 mice infected with Mtb Erdman were pooled at 21 days post-infection. IFNγ and IL-17 concentrations in the cell culture supernatant revealed responses targeted towards amino acids 92-116 and 172-203. Graphs represent one experiment with one sample per 3 peptide pool. D) Peptide binding affinity for the H2-IAb allele was predicted using IEDB NetMHCIIpan 4.0 BA using the RimJ sequence and 15-18 amino acid peptide length. Predicted binding affinity ranked by percentile ranking was graphed for the best predicted binding peptides for the designated regions of RimJ. The prediction revealed similar hotspots to the regions identified in the stimulation experiments (A-C). Labeled rectangles (99-116 and 182-197) are the peptides predicted to be stimulate the majority of the response to RimJ-Erdman.

**Supplemental Figure 3.**
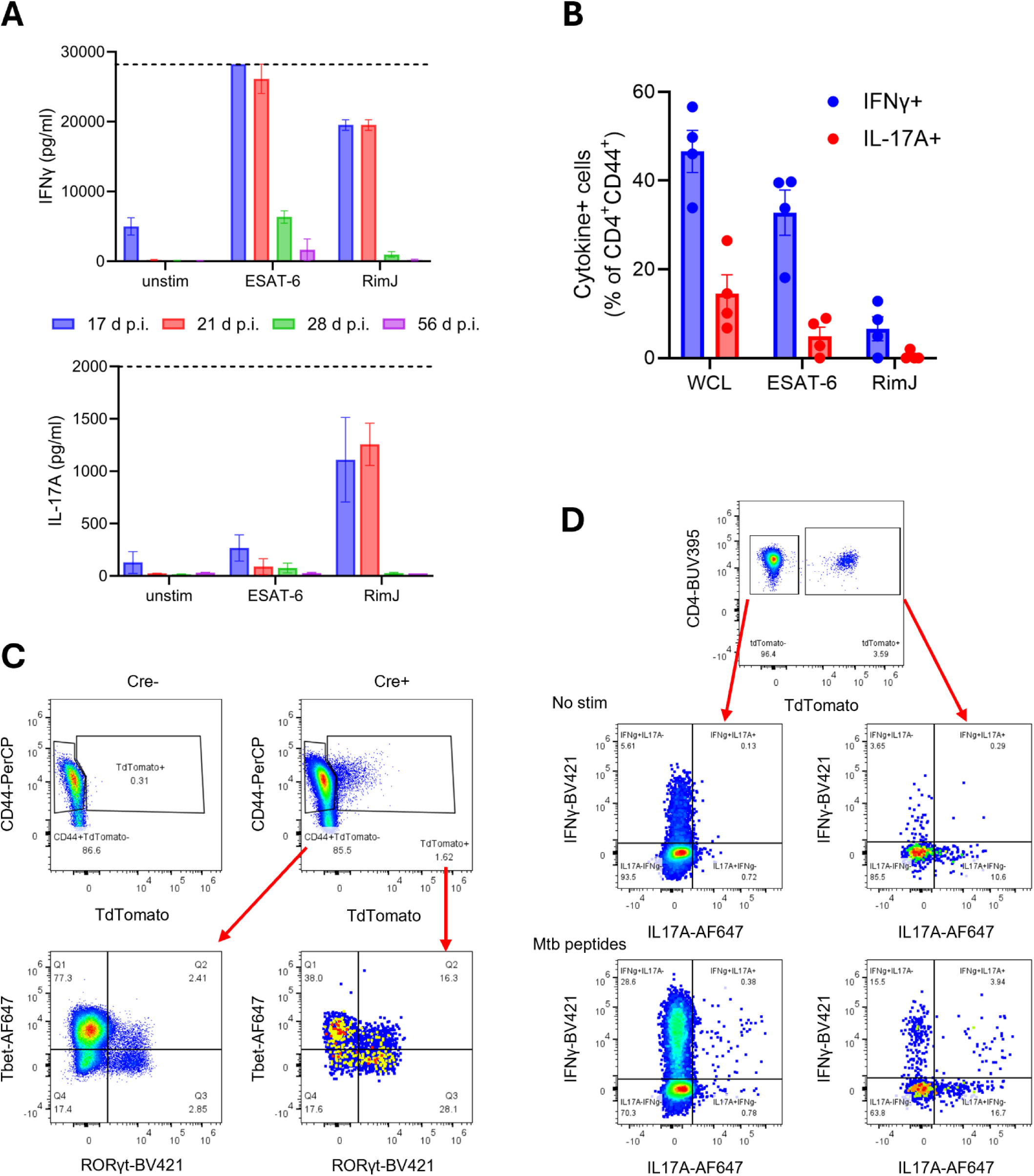
IL-17A response to RimJ in the mediastinal lymph node and IL-17A reporter expression in CD4 T cells during Mtb infection. A) C57BL/6 mice were infected with Mtb Erdman, and mediastinal lymph nodes were pooled from 5 mice at each timepoint and cultured in triplicate in the presence of ESAT-6 peptides (10 μg/ml), RimJ peptides (10 μg/ml), or no peptides for 72 hours. IFNγ and IL-17A were measured in the culture supernatants by ELISA. B) Lung cells from C57BL/6 mice infected with Mtb Erdman 28 days prior, were stimulated with Mtb whole cell lysate (10 μg/ml), ESAT-6 peptides (10 μg/ml), or RimJ peptides (10 μg/ml) for ICS. Frequencies of IFNγ^+^ or IL-17A^+^ CD4 T cells were determined by flow cytometry. C-D) *Il17a-cre*; Ai14 reporter mice were infected with Mtb Erdman and used to identify IL-17A-producing cells by fate-mapping. C) Transcription factor staining of lung cells from infected reporter mice. D) Stimulation of lung cells for ICS to identify IFNγ^+^ and IL-17A^+^ cells.

**Supplemental Figure 4.**
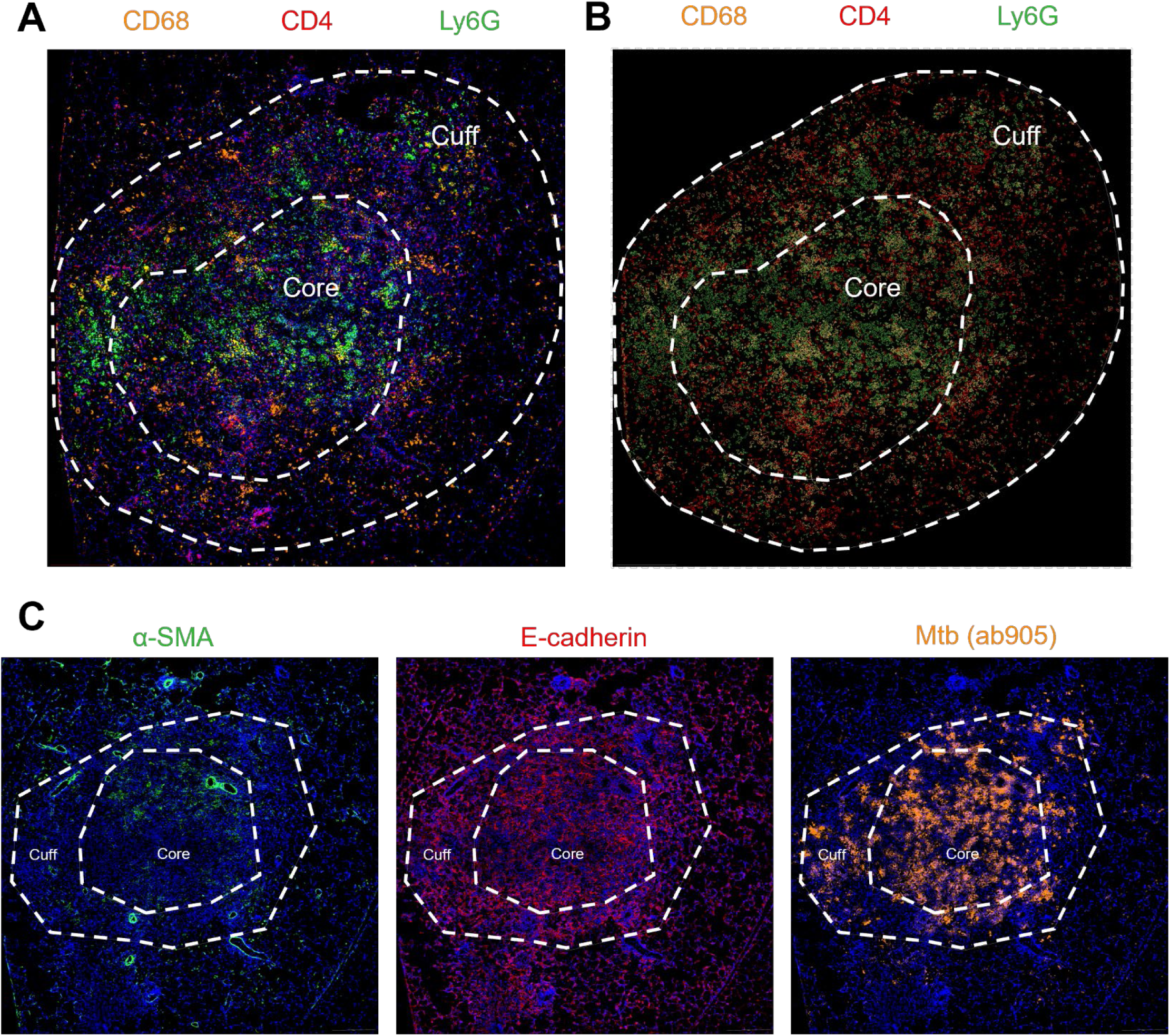
Analysis of lung immunofluorescence images. A) Representative images depicting the manual assignment of the core and cuff regions of the identified lesions. B) Masking of the positive signal for each marker using pixel thresholders in QuPath to create objects for CD4^+^ and CD68^+^ cells and measure total area for Ly6G, α-SMA, E-cadherin, and Mtb (ab905) staining. C) Dual channel images showing α-SMA (green), E-cadherin (red), and Mtb (ab905, orange) with DAPI counterstain from a representative pCas4-vaccinated mouse lesion.

**Supplemental Figure S5.**
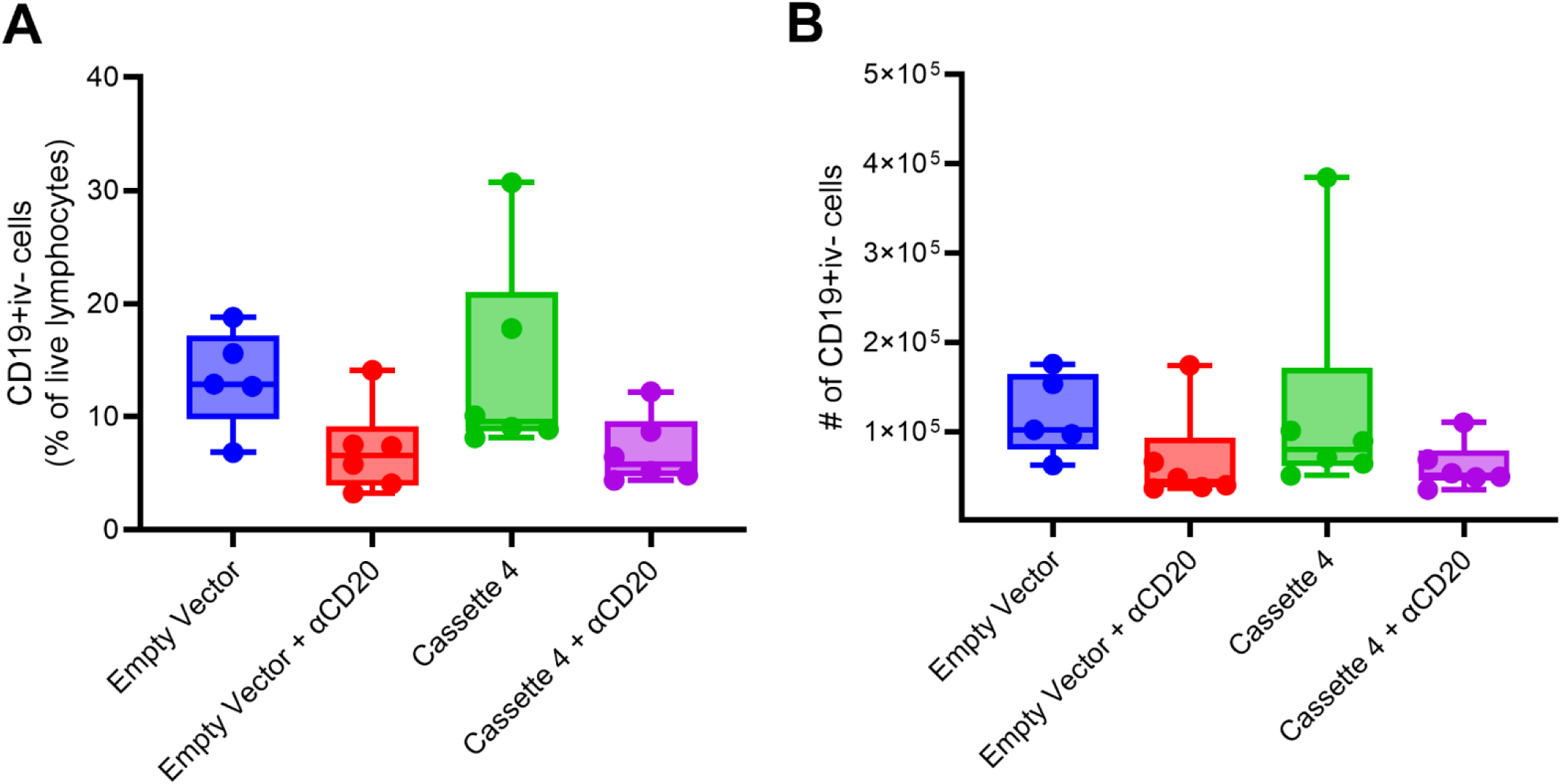
B cell numbers in the lungs of SP140^-/-^ mice at 21 days post-infection after anti-CD20 B cell depletion during vaccination. Mice vaccinated with mock (Empty Vector) or pCas4 (Cassette 4) and treated with anti-CD20 antibody or not were challenged with Mtb Erdman and lung cells were evaluated by flow cytometry at 21 days post-infection. The frequency (A) and total number (B) of parenchymal B cells (CD19^+^ cells) was slightly reduced in mice treated with anti-CD20 antibody, but not significantly different.

